# Epigenetic modifications mediate cultivar-specific root development and metabolic adaptation to nitrogen availability in wheat

**DOI:** 10.1101/2023.02.24.529862

**Authors:** Hao Zhang, Fa Cui, Long Zhao, Xiaoyu Zhang, Jinchao Chen, Jing Zhang, Yongpeng Li, Yanxiao Niu, Lei Wang, Caixia Gao, Xiangdong Fu, Yiping Tong, Hong-Qing Ling, Junming Li, Jun Xiao

**Affiliations:** State Key Laboratory of Plant Cell and Chromosome Engineering, Institute of Genetics and Developmental Biology, Chinese Academy of Sciences, Beijing 100101, China; Key Laboratory of Molecular Module-Based Breeding of High Yield and Abiotic Resistant Plants in Universities of Shandong, College of Agriculture, Ludong University, Yantai 264025, China; University of Chinese Academy of Sciences, Beijing 100049, China; Center for Agricultural Resources Research, Institute of Genetics and Developmental Biology, Chinese Academy of Sciences, Shijiazhuang 050022, Hebei, China; Ministry of Education Key Laboratory of Molecular and Cellular Biology, Hebei Research Center of the Basic Discipline of Cell Biology, Hebei Key Laboratory of Molecular and Cellular Biology, College of Life Sciences, Hebei Normal University, Shijiazhuang 050024, China; Hebei Collaboration Innovation Center for Cell Signaling, Shijiazhuang, 050024, China; Hainan Yazhou Bay Seed Laboratory, Sanya, Hainan, China; Centre of Excellence for Plant and Microbial Science (CEPAMS), JIC-CAS, Beijing, China

**Keywords:** epigenetic regulation, root architecture, nitrogen uptake, wheat

## Abstract

The breeding of crops with improved nitrogen use efficiency (NUE) is crucial for sustainable agriculture. However, the role of epigenetic modification in the regulation of cultivar-specific responses to low nitrogen (LN) constraint is not well understood. Here, we analyzed the chromatin landscapes in roots, leaves, and seeds in two wheat cultivars (KN9204 and J411) that differ radically in NUE under normal and nitrogen-limited conditions. Transcriptional regulatory chromatin regions exhibited a clear cultivar-specificity between KN9204 and J411. Cultivar-specific regulation of nitrogen metabolism genes (NMGs) is linked to variation in histone modification levels, rather than differences in DNA sequence. Cultivar-specific histone modification regions were found to contribute to the genetic regulation of NUE-related traits, such as QTL locus of maximum root length of qMRL-7B. Furthermore, LN-induced H3K27ac and H3K27me3 dynamics enhanced root growth more in KN9204, while strengthened the nitrogen uptake system more in J411. Evidence from histone deacetylase inhibitor treatment and transgenic plants with loss-of-function of the H3K27me3 methyltransferase further showed that changes in epigenetic modifications can alter the strategy for root development and nitrogen uptake in response to LN constraint. Taken together, the results of our study highlight the importance of epigenetic regulation in mediating cultivar-specific root development and metabolic adaptation to LN in wheat.

## Introduction

Enhancing nitrogen-use efficiency (NUE) in crops is a critical requirement in modern agriculture to increase yields and maintain sustainability (Chen et al., 2014). NUE is determined by two components; N uptake efficiency (NUpE), i.e., nitrogen acquisition from the soil, and N utilization efficiency (NUtE), i.e., yield per unit of nitrogen acquired. NUE is affected by nitrogen uptake, assimilation, remobilization, and storage (Masclaux-Daubresse et al., 2010; Wang et al., 2018; Xiao et al., 2022). Various agronomic traits have been studied as indexes to uncover the mechanism for NUE regulation, such as root morphology (Forde, 2014; Kiba and Krapp, 2016), tissue-specific nitrogen contents (Hu et al., 2015; Li et al., 2017), and plant architectures including tiller number and plant height (Liu et al., 2021; Sun et al., 2014). Several regulators have been identified for mediating crop growth under nitrogen-limited conditions. For example, Teosinte branched1, Cycloidea, Proliferating cell factor (TCP)-domain protein 9 (*OsTCP19*) was reported to modulate tillering in rice in response to different nitrogen availability (Liu et al., 2021). The ‘Green Revolution’ gene Reduced height-1 (*Rht1*) reduces root size, while introduction of the 1 RS alien chromosome translocation from rye greatly increased shallow and deep root growth and thus N uptake in wheat (Waines and Ehdaie, 2007). Manipulating factors in the nitrogen metabolism process could improve NUE in crops; examples are efficient nitrogen uptake by an elite allele of the transporter gene *NRT1.1B* (Hu et al., 2015), *NRT1.7* (Chen et al., 2020), and nitrate reductase *NR2* (Gao et al., 2019), as well as the assimilation factors *GS* (Li et al., 2011), and *GOGAT* (Quraishi et al., 2011). In addition, changes to the transcriptional regulation of the nitrogen metabolic process is also important for improving nitrogen use (Gaudinier et al., 2018). Overexpression of *TaNAC2-5A* (NAM, ATAF, and CUC) increased the nitrate influx rate, N uptake, and grain N concentration by positively regulating *TaNRT2.5* and *TaNRT2.1* expression (He et al., 2015). TabZIP60, a basic leucine zipper transcription factor, can bind and negatively regulate *TaGOGAT-3B* expression, which affects N uptake and grain yield (Yang et al., 2019).

Recent studies have highlighted the importance of epigenetic regulation in modulating nitrogen metabolism and the plant response to nutrient availability (Sere and Martin, 2020). SET DOMAIN GROUP 8 (SDG8) regulates the N assimilation and lateral root response by regulating H3K36me3 level in response to changing N levels in *Arabidopsis* (Li et al., 2020). HIGH NITROGEN INSENSITIVE9 (*AtHNI9*) regulates root nitrate uptake by affecting the H3K27me3 level at the *AtNRT2.1* locus (Widiez et al., 2011). In rice, NITROGEN-MEDIATED TILLER GROWTH RESPONSE 5 (*NGR5*), an APETALA2-domain transcription factor, recruits the Polycomb repressive complex 2 (PRC2) to deposit H3K27me3 and repress the expression of branching-inhibitors such as the strigolactone signaling gene *Dwarf14* (*D14*) and *squamosa promoter binding proteinlike 14* (*OsSPL14*) (Wu et al., 2020). This regulatory module enables nitrogen-dependent stimulation of rice tillering (Wu et al., 2020). However, the role of epigenetic regulation in the nitrogen-use process in wheat is largely unexplored.

Kenong 9204 (KN9204) and Jing 411(J411) are represent cultivars grown in the North China Plain with diverse agriculture traits (root architecture and nitrogen uptake) under low nitrogen constraint (Wang et al., 2011). Recombinant inbred lines (RILs) generated between cross of KN9204 and J411 were widely used for localization of NUE-related QTL traits (Cui et al., 2014), such as root length, root tips, and grain protein content by us and others (Cui et al., 2016; Fan et al., 2018; Zhao et al., 2018). We recently completed the reference genome sequence of KN9204 and identified 882 nitrogen metabolism genes (NMGs) (Shi et al., 2022). Comparative transcriptome analysis between KN9204 and J411 revealed different responsive programs under low nitrogen constraint, especially in the yield-related spike tissue and in seeds during reproductive development (Shi et al., 2022). The underground tissues (roots) are not as well investigated, and the regulatory mechanisms that control transcriptional program diversity and how the differences are correlated with varied NUE in different cultivars are unclear.

In this study we profiled epigenomic modifications in KN9204 and J411 via CUT&Tag (Kaya-Okur et al., 2019) for various histone modifications (H3K27ac, H3K27me3, H3K4me3, H3K36me3, H3K9me3) and the histone variant H2A.Z, in three tissues under different nitrogen conditions. The epigenomic maps provide a systematic view of the dynamic chromatin landscapes of two cultivars that differ in NUE in response to LN constraint. Integrating the data with previously generated QTL analysis uncovered insights into cultivar-specific transcription-bias and epigenomic divergence, especially H3K27ac, in response to LN constraint. Changes to H3K27ac and H3K27me3 by chemical inhibitors or genetic mutations provide further evidence that manipulating histone modification profiles can adjust the LN adaptation strategy selection in different wheat cultivars.

## Results

### Profiling the tissue-specific chromatin landscape of wheat cultivars under different nitrogen conditions

To understand the epigenetic regulation of the transcriptomic dynamics of wheat cultivars that differ in NUE in response to limited nitrogen (N) supply, we did CUT&Tag of various histone modifications for the wheat cultivars KN9204 and J411 at normal nitrogen (NN) and LN conditions for roots (28 day), flag leaves (heading stage) and seeds (21 days after anthesis) (Fig. 1a), corresponding to transcriptomes generated previously (Shi et al., 2022).

**Fig. 1.**
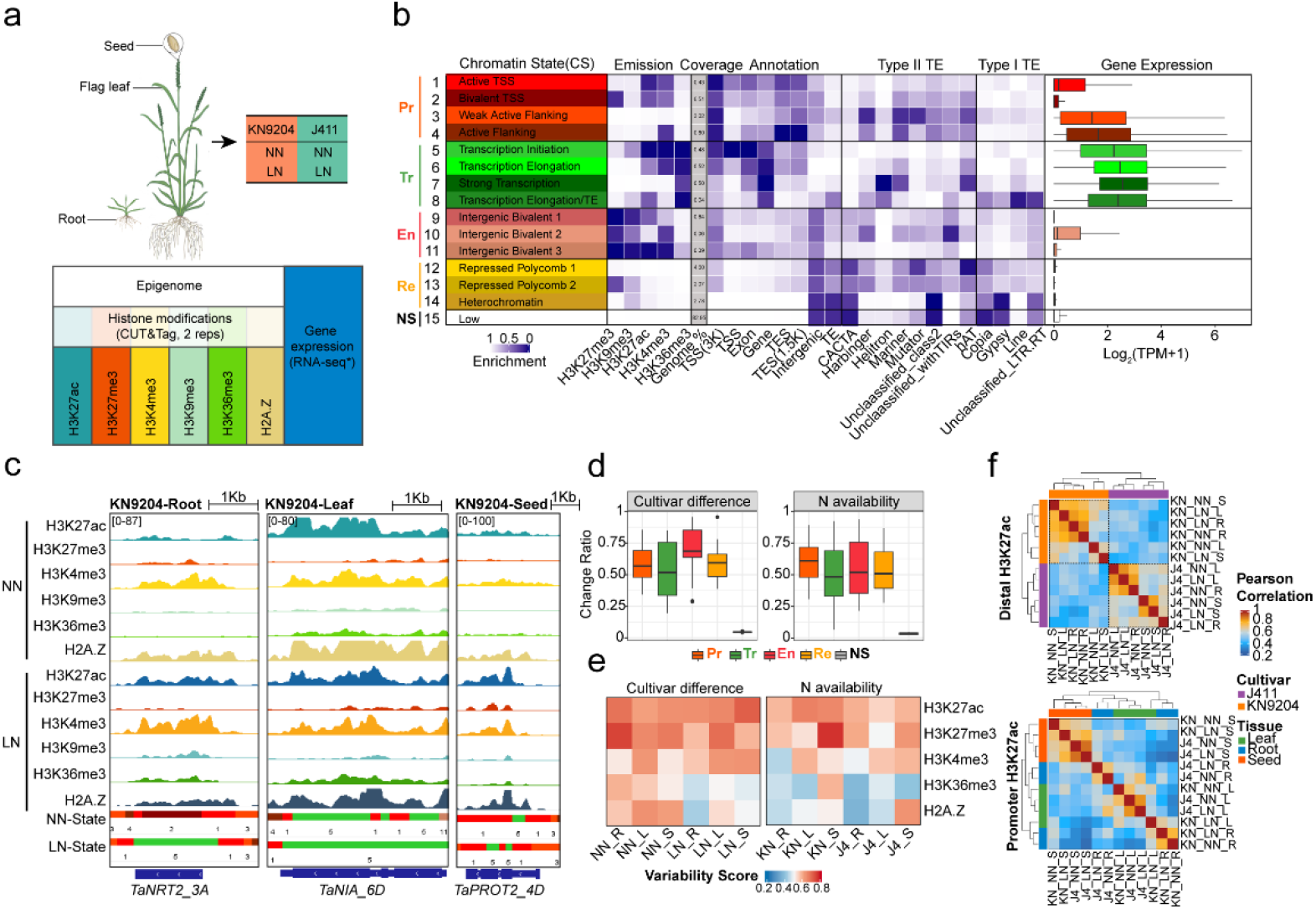
A 15-state model characterizes the dynamic chromatin landscape in KN9204 and J411. a. Experimental design for generating the epigenomic data sets in KN9204 and J411 under different nitrogen conditions. b. Chromatin state definitions, genomic annotation enrichments, and expression levels of genes associated with each chromatin state. Chromatin states could be mainly organized into five categories: Pr (promoter), Tr (transcription), En (enhancer-like), Re (repressive), and Ns (No signal) states. c. Schematic diagrams representing the chromatin state dynamics under different nitrogen conditions and tissues. State colors are as in (b). d. The fractions of bases for the states of the five categories that vary between cultivar difference and nitrogen availability. e. Variability scores of histone modifications between cultivar difference and nitrogen availability. f. Cross-correlation heatmaps of all H3K27ac peaks which were located in the distal or promoter regions separately for the two cultivars and the two nitrogen conditions.

The root transcriptome is more sensitive to LN treatment as compared to flag leaves or seeds, with more differentially-expressed genes (DEGs) for both KN9204 and J411 under LN/NN conditions (Fig. S1a). This is consistent with phenotypic variations in the different tissues in KN9204 and J411, as we previously reported (Shi et al., 2022). In roots, genes associated with hormone-related processes, such as strigolactone biosynthesis and the auxin efflux transmembrane transporter, are induced by LN in KN9204, while expression of the high-affinity transporter of nitrite (*NRT*) genes was highly induced in J411 under LN (Fig. S1b). In flag leaves and seeds, sugar metabolism and protein storage processes were activated by LN in KN9204, while stress-related processes were induced in J411 by LN (Fig. S1b). Thus, wheat cultivars that differ in NUE have different transcriptional program dynamics in response to LN.

The CUT&Tag assay showed reproducibility between two biological replicates for the various histone modifications (Fig.S1c). The majority of histone modification peaks were located in the distal region expected for H3K36me3 in wheat (Fig. S1d), as described in our recent report (Zhao et al., 2023). H3K27ac and H3K4me3 were associated with highly expressed genes, while H3K27me3 is enriched in low/non-expressed genes (Fig. S1e). H3K4me3 and H2A.Z did not show a preference for gene expression level (Fig. S1e). H3K9me3 was associated with transposable elements (TEs), because more than half of the peaks were located in TE regions (Fig.S1f). Genes involved in biotic and abiotic stress responses had a higher H2A.Z level relative to that in developmental and hormone-related genes (Fig.S1g). Furthermore, we used ChromHMM (Ernst and Kellis, 2017) to systematically examine the chromatin state defined by combinatorial patterns of various histone marks (Fig. 1b). The whole genome could be partitioned into fifteen chromatin states (CS) forming five major categories; Promoter (CS1-4), Transcription (CS5-8), Enhancer like (CS9-11), Repressive (CS12-14), and No signal (CS15), with different genome coverage, TE enrichment, and gene expression level (Fig.1b). The Promoter and Enhancer-like states were both associated with H3K27ac, H3K4me3, and H3K27me3 but were located in the transcription start site (TSS) or the intergenic region, respectively (Fig. 1b). Repressive states are mainly enriched with H3K27me3, covering about 10% of the genome, whereas, no signal state accounted for a major part (∼83%) of the genome. Thus, a very limited part of the wheat genome (∼7%) is transcribed or involved in transcriptional regulation in the context of various histone modifications (Fig. 1b).

Next, we asked how chromatin state dynamics reflect the differences between wheat cultivars, tissue types, and nitrogen conditions. For example, genes related to nitrate uptake in the root (*TaNRT2_3A*), assimilation in the leaf (*TaNIA_6D*), and amino acid storage in the seed (*TaPROT2_4D*) showed varied CS under LN condition in KN9204 (Fig. 1c). Generally, the Enhancer-like CS was the most variable chromatin region between the two different cultivars, whereas the Promoter CS is mostly affected by nitrogen availability (Fig. 1d). CS10 and CS9 of the Enhancer-like states showed a higher frequency of change between the wheat cultivars, and CS10 and CS2 varied more under different nitrogen availabilities (Fig. S1h). Furthermore, we calculated variability scores to estimate which histone marks might mediate chromatin state change (Fig. 1e). H3K27ac and H3K27me3 were at top of the variability score as compared to others for the different nitrogen conditions and between the wheat cultivars (Fig. 1e). Based on the distribution pattern (Fig. S1d), we subdivided the peaks of these two histone marks into distal and promoter categories and evaluated their variability using Pearson correlation analysis (Fig.1f, Fig.S1i). Distal H3K27ac showed a clear cultivar-specificity, whereas promoter H3K27ac showed more tissue-specificity (Fig. 1f). For H3K27me3, both distal and promoter peaks showed clear cultivar-specificities (Fig. S1i).

Therefore, wheat cultivars, tissue types and nitrogen conditions shape distinct genomic regions with variable chromatin states, especially for H3K27ac and H3K27me3.

### Cultivar-biased expression of nitrogen metabolism genes is mainly mediated by variations in histone modifications

Nitrogen metabolism genes (NMGs) are crucial for plants to take up and utilize nitrogen, and these include genes that encode nitrate transporter (NPF, NRT2, NAR2, CLC, and SLAH), nitrate reductase (NIA, NIR), ammonium transporter (AMT), and amino acid transporter (APC) family members, as well as genes for proteins involved in ammonium assimilation (GS, GOGAT, GDH, ASN, AspAT) and transcription factors (TFs) related to N metabolism (Krapp, 2015; Wang et al., 2018). Previously, we have identified 882 NMGs in wheat by sequence similarity comparisons to six plant species (*Brachypodium distachyon*, barley, rice, sorghum, maize, and *Arabidopsis thaliana*) (Shi et al., 2022).

We compared the NMGs between KN9204 and J411 for DNA sequence variation at H3K27ac-marked promoter regions, transcriptional level changes, and changes in histone modifications. In general, approximately 25% of NMGs showed different expression levels between KN9204 and J411 (FDR<0.05, fold-change≥1.5), while only about 5% of NMGs have DNA sequence variations in the promoter regulatory regions (Fig. 2a). The variation in expression of NMGs between KN9204 and J411 existed for different gene families in various tissues (Fig. 2b). In particular, the expression levels of *NRT2* and *NIA* in J411 were both higher under the LN condition relative to KN9204 in roots, whereas expression of *NPF* family genes was activated in KN9204 roots (Fig. S2a). In flag leaves, *NIA* genes were up-regulated in KN9204 but not in J411. As for seeds, the relative expression of genes that encode NPF and GS was much higher in KN9204 compared to J411 (Fig. S2a).

**Fig. 2.**
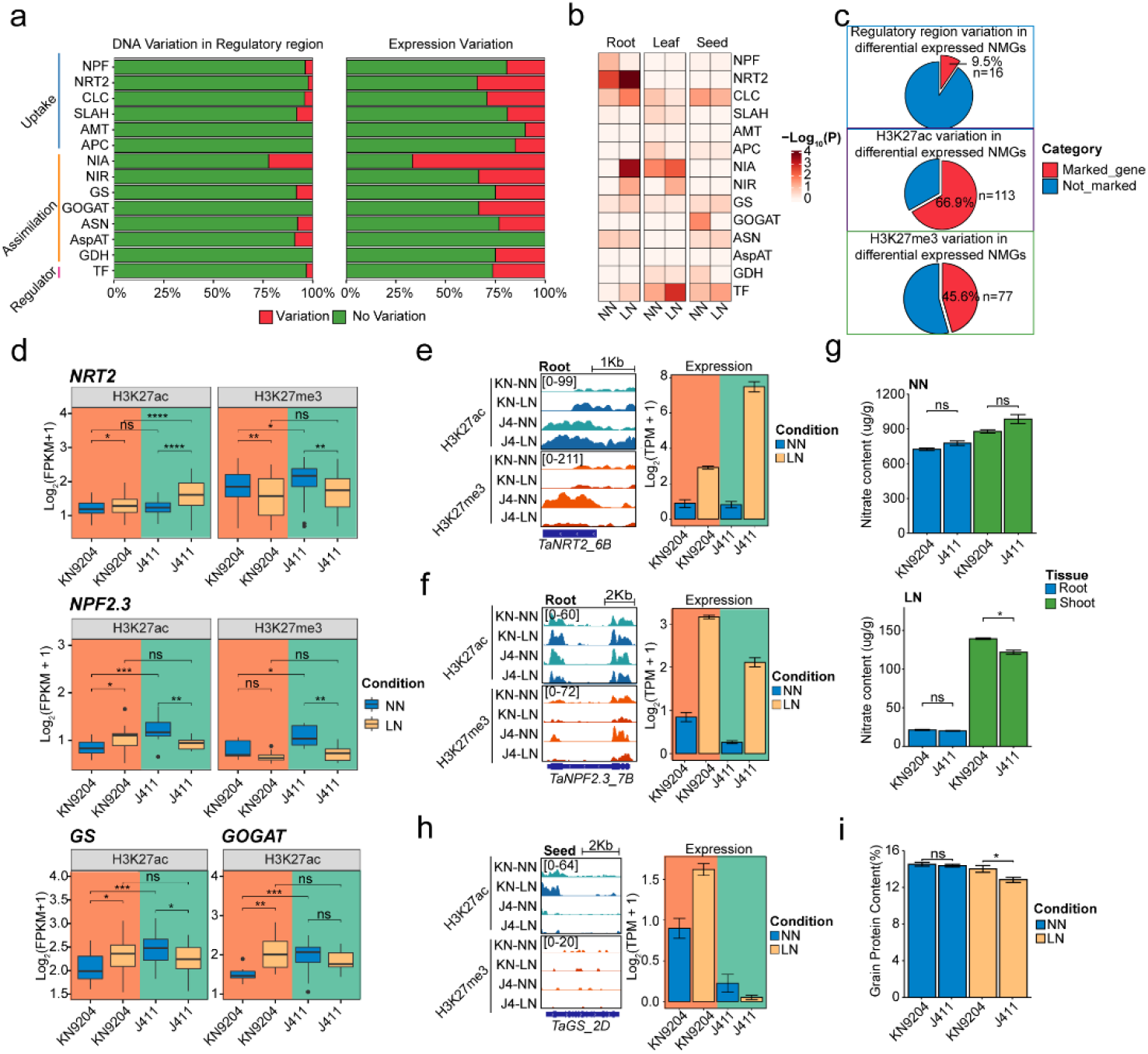
Epigenomic variations are vital to the expression bias of NMGs. a. Fraction of the regulatory regions and expression variation of nitrogen metabolism genes (NMGs) between wheat cultivars KN9204 and J411. b. Enrichment of LN-induced differentially-expressed NMGs based on their functional category in three tissues (Fisher’s exact test was used to calculate the *p*-values for the overlaps). c. Proportion of NMGs showing expression variation between KN9204 and J411 with regulatory region variation (top panel), H3K27ac variation (middle panel), and H3K27me3 variation (bottom panel). d. Histone modification (H3K27ac and H3K27me3) levels of NRT2 and NPF2.3 in the roots of KN9204 and J411 between the two nitrogen availability levels, and dynamic H3K27ac levels of GS and GOGAT in the seeds (Wilcox test: ns: p>0.05; *: *p*≤0.05; **: *p*≤0.01; ***: *p*≤0.001; ****: *p*≤0.0001). e. Representative tracks showing histone modifications and transcriptional changes of *TaNRT2_6B* in roots for the two wheat cultivars and two nitrogen levels. f. Representative tracks showing histone modifications and transcriptional changes of *TaNPF2.3_7B* in roots for the two wheat cultivars and two nitrogen levels. g. Nitrate content of shoots and roots of KN9204 and J411 under different nitrogen conditions (Student’s *t*-test; ns: *p*>0.05; *: *p*≤0.05). h. Representative tracks showing histone modifications and transcriptional changes of *TaGS_2D* in seeds for the two wheat cultivars and two nitrogen levels. i. Grain protein content (GPC) in seeds of KN9204 and J411 under different nitrogen conditions (Student’s *t*-test; ns: *p*>0.05; *: *p*≤0.05).

Cultivar-biased expression of NMGs is not likely to be caused by variation in DNA sequence in the regulatory regions between the two cultivars, because only 16 genes overlapped (Fig. 2c). Instead, we found that the majority of NMGs that showed cultivar-biased expression were marked by differential peaks of H3K27ac (66.9%) and H3K27me3 (45.6%) (Fig. 2c; Fig. S2b, Supplemental Table S1). In roots, the *NRT2* family showed similar trends for changes in H3K27ac and H3K27me3 under LN/NN conditions in both cultivars, but the changes were more dramatic in J411 compared to KN9204 (Fig. 2d, top). For example, at the *TaNRT2_6A* (*TraesCS6A02G031000*) locus, H3K27ac gains and H3K27me3 losses in response to LN occurred in both cultivars but were more dramatic in J411 (Fig. 2e). This fits well with the higher level of induced expression of *TaNRT2_6A* in J411 compared to KN9204 under LN conditions (Fig. 2e). Expression of *TaNPF2.3*, which is involved in nitrite transport from the root to the shoot (Taochy et al., 2015), was activated in KN9204 accompanied by a decrease H3K27me3 under LN (Fig. 2d, middle). However, no change in H3K27me3 but a significant decrease in H3K27ac was observed in J411 (Fig. 2d, middle). *TaNPF2.3_7B* expression was increased in KN9204 but decreased in J411 under LN/NN conditions (Fig. 2f). We found that the nitrate contents in shoots relative to roots were consistently higher in KN9204 compared to J411 under LN conditions (Fig. 2g). In seeds, for the ammonium assimilation family genes for GS and GOGAT, the trends for H3K27ac abundance in response to LN were opposite in the two cultivars; relative expression increased in KN9204 and decreased in J411(Fig. 2d bottom). The expression of *TaGS_2D* increased in KN9204 but decreased in J411 under LN/NN conditions, which was accompanied by an increase in the H3K27ac level in KN9204 (Fig. 2h). Coincidently, KN9204 seeds have a higher protein content compared to J411 under LN conditions (Fig. 2i).

Taken together, the varied levels of H3K27ac and H3K27me3 are associated with the expression bias of NMGs, and this then results in divergent nitrogen metabolism processes in KN9204 and J411.

### Cultivar-specific H3K27ac regions influence NUE-related traits by divergent transcriptional regulation

Given the importance of H3K27ac in mediating chromatin status changes and the expression dynamics of NMGs between cultivars and different tissues at LN/NN, we further identified cultivar-specific H3K27ac peaks by K-means clustering (Fig. 3a). In general, the cultivar-specific H3K27ac peaks are over-represented in distal regions and under-represented in promoter and genic regions as compared to all H3K27ac peaks (Fig. 3b). Cultivar-specific H3K27ac-marked promoters are associated with cultivar-specific expressed genes in roots, leaves, and seeds (Fig. 3c, Fig. S3a, Fig. S3b). For example, KN9204-specific H3K27ac-marked genes are associated with nutrient reservoir activity and cell wall biogenesis (e.g., *PROT1, LBD16, XTH19, CSLC5*) (Fig. 3c, Fig. S3c), whereas genes involved in flavonol biosynthesis are specifically marked by H3K27ac in J411 (Fig. S3c).

**Fig. 3.**
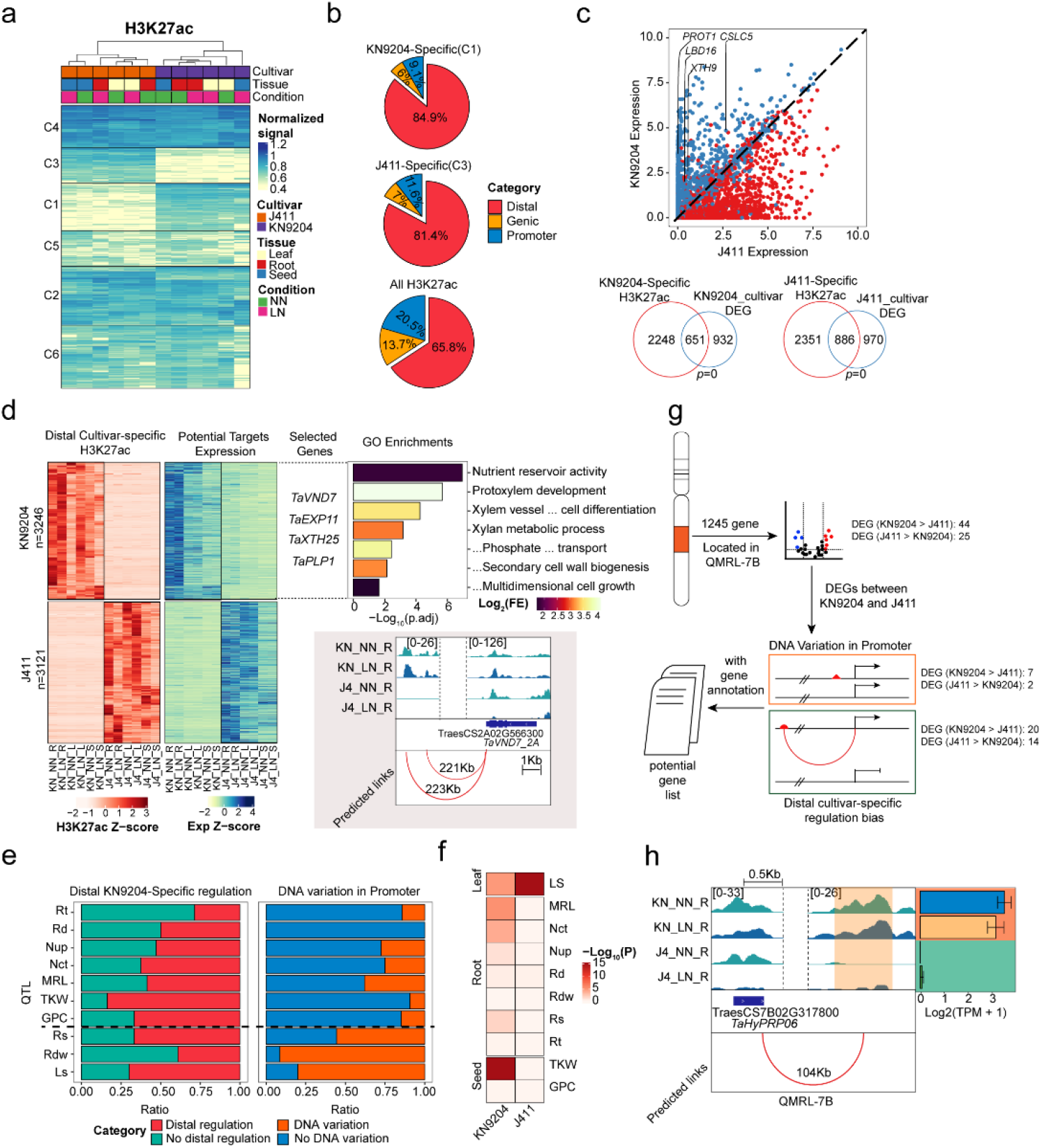
Distal regulatory region divergence is involved in NUE-trait variation between KN9204 and J411. a. K-means clustering of the variable H3K27ac peaks. b. Peak distribution (distal, genic, or promoter) of cultivar-specific H3K27ac peaks in the wheat genome. c. Mean gene expression in J411 (x-axis) versus mean gene expression in KN9204 (y-axis) for genes associated with proximal cultivar-biased H3K27ac peaks separately in the roots (top panel). Overlap between genes marked by cultivar-specific H3K27ac and DEGs between cultivars (blue circle) (bottom panel) (Fisher’s exact test was used to calculate the *p*-values for the overlaps). d. Assignment of distal cultivar-specific H3K27ac peaks to potential targets; representative genes and GO enrichment are shown on the right, and representative case is shown at the bottom. e. Fraction of DEGs with distal KN9204-specific regulation (left) or DNA variation within the promoters located in QTLs (right) between KN9204 and J411. Rt: root tip number; Rd: root diameter; Nup: Nitrogen uptake content; Nct: Nitrogen concentration; MRL: Maximum root length; TKW: Thousand-kernel weight; GPC: Grain protein content; Rs: Root surface area; Rdw: Root dry weight; Ls: flag leaf size. f. Enrichment of target genes with distal cultivar-specific H3K27ac peaks in QTLs between KN9204 and J411 (Fisher’s exact test was used to calculate the *p*-values for the enrichment). Abbreviations as in (e). g. Schematic diagram illustrating the process used to narrow down potential key genes regulated by distal cultivar-specific regulation in the QTL regions (*QMRL-7B* in this case). h. Representative tracks of *TaHyPRP06* regulated by distal cultivar-specific H3K27ac peaks in *QMRL-7B*.

The cultivar-specific H3K27ac peaks in distal region are co-localized with cultivar-specific H3K4me3 and H2A.Z (Fig. S3d), indicating that such distal epigenetically modified hotspots may function as ‘enhancers’ in regulating gene expression (Calo and Wysocka, 2013; Giaimo et al., 2019; Russ et al., 2017). We further assigned those regions to potential targets within 500 kb distance by correlation between H3K27ac and genes expression dynamics as reported previously (Fig. S3e)(Wang et al., 2021). In total, 58,493 unique pairs (22,003 distal H3K27ac regions and 6,357 target genes) were identified between the distal cultivar-specific H3K27ac peaks and genes (Fig. 3d). The accuracy of distal H3K27ac-to-gene pairs was supported by higher chromatin interaction ratio at the anchors from published Hi-C data (Fig. S3f). GO annotation of KN9204-specific distal H3K27ac-regulated genes showed enrichment of genes for cell wall synthesis, protoxylem development, and nutrient reservoir activity, which is similar to the KN9204-specific promoter H3K27ac-marked genes. Among them, several genes have been reported to function in nutrient transport to the shoot, such as *TaVND7* (Yamaguchi et al., 2011), *TaXTH25* (Sasidharan et al., 2010). No GO category was enriched in targets of J411-specific distal H3K27ac (Fig. 3d).

We analyzed the contribution of cultivar-specific regulatory regions to the DEGs located within QTLs identified via linkage analysis in a KN9204-J411 RIL population (Cui et al., 2016; Fan et al., 2018; Zhao et al., 2018). Indeed, compared with DNA sequence variations in promoter regions, more cultivar-specific distal H3K27ac regions are correlated with differentially-expressed genes within QTLs regions such as nitrogen uptake (Nup), nitrogen concentration (Nct), maximum root length (MRL), grain protein contents (GPC) (Fig. 3e, Fig. S3g). The target genes of KN9204-specific H3K27ac peaks were significantly enriched in those QTLs of the NUE related-traits, in which the elite genetic loci originated from KN9204 (Fig. 3f, supplemental table S2). The J411-specific H3K27ac region associated genes were enriched within leaf size-related QTL (Fig. 3f). Taking the previously identified qMRL-7B as an example (Fan et al., 2018), 1,245 genes are located within the genetic region with 69 DEGs and only nine genes contained DNA sequence variations within the promoter regions, while 34 genes had cultivar-specific distal H3K27ac peaks (Fig. 3g). Among them, *TraesCS7B02G317800* (*TaHyPRP06_6B*) and *TraesCS7B02G326900* (*TaXTH25_7B*) have been reported to regulate root development (Zhao et al., 2022b), and the higher expression levels in KN9204 compared to J411 were correlated with KN9204-specific H3K27ac distal regulatory regions (Fig.3h; Fig. S3h).

Thus, both promoter and distal cultivar-specific H3K27ac regions are associated with transcriptional regulation and contains genetic variations that regulate NUE-related traits.

### LN-induced H3K27ac dynamics affect divergent adaptive programs in KN9204 and J411

WE next wondered how LN-induced H3K27ac dynamics could trigger divergent adaptation in the cultivars KN9204 and J411. H3K27ac shows varied dynamic patterns in roots, flag leaves, and seeds in response to LN in KN9204 and J411 (Fig. 4a). In roots, H3K27ac showed subtle changes in KN9204, while drastic loss and relatively less gain were apparent in J411 under LN/NN conditions. K-means clustering further identified different categories of dynamic H3K27ac regions in roots, flag leaves, and seeds, under LN conditions for KN9204 and J411 (Fig. 4b, Fig. S4a).

**Fig. 4.**
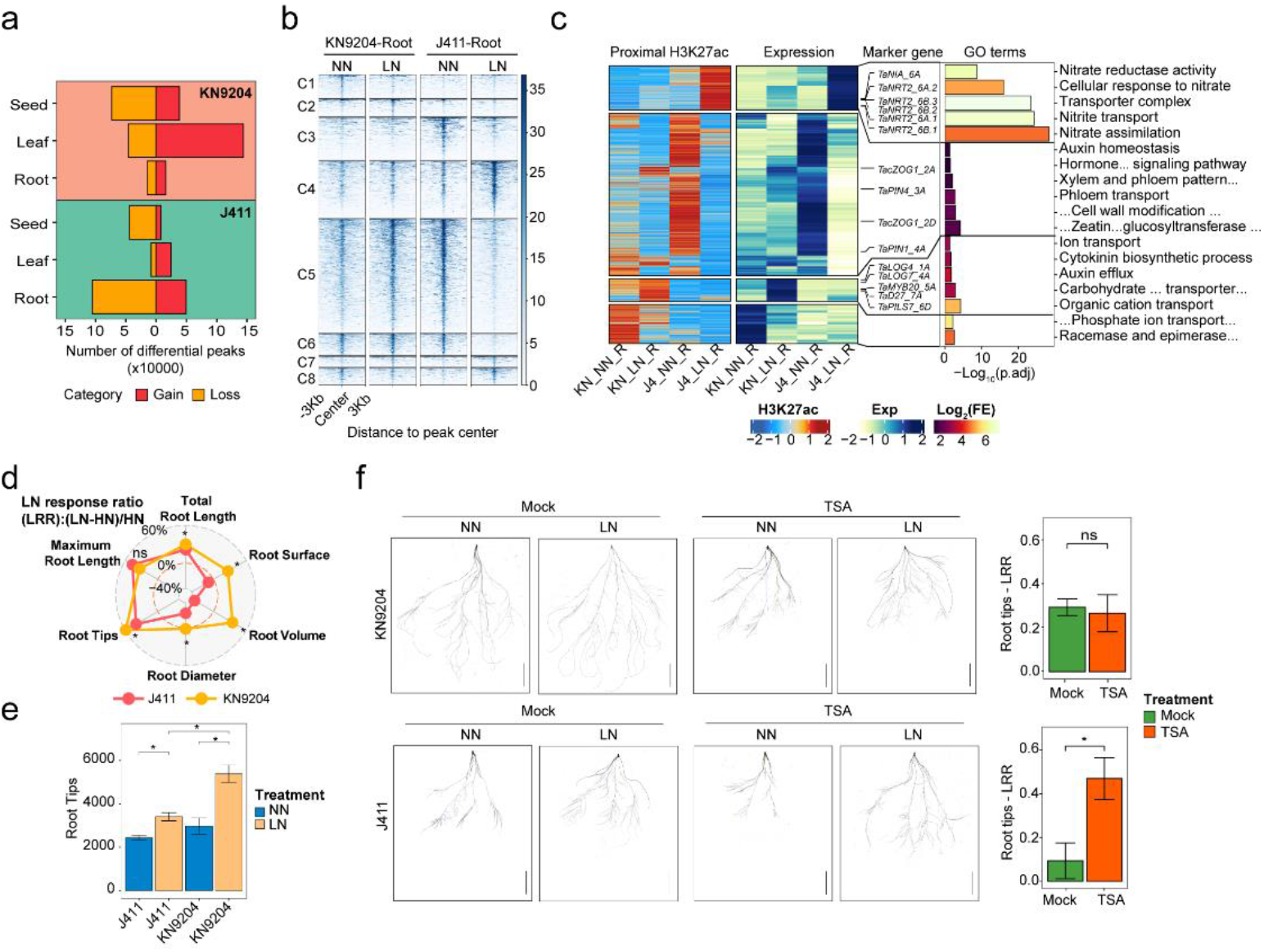
Dynamics of H3K27ac reveals the divergent strategies of KN9204 and J411 to low nitrogen conditions. a. The number of differential H3K27ac peaks in response to LN in three tissues (seeds, roots, leaves) in KN9204 and J411. b. Heatmaps showing the differential H3K27ac peaks in eight clusters in the roots of KN9204 and J411 under NN and LN conditions. c. Dynamic proximal H3K27ac peaks and corresponding gene expression changes in the roots of KN9204 and J411. Representative genes and GO (gene ontology) enrichments are shown on the right. d. The LN-response-ratio (LRR) of root systems for KN9204 and J411 (Student’s *t*-test; ns: *p*>0.05; *: *p*≤0.05). e. The number of root tips in KN9204 and J411 plants under different nitrogen conditions (Student’s *t*-test; ns: *p*>0.05; *: *p*≤0.05). f. Phenotypes of the root systems of KN9204 and J411 seedlings in the control (mock) and TSA treatments (2 μM) under different nitrogen conditions. Scale bars=5cm. The right panel is the LRR of the root tips (Student’s *t*-test; ns: *p*>0.05; *: *p*≤0.05).

Extensive loss of H3K27ac was observed in roots of J411 compared to KN9204 (clusters C3, C5, and C6, n=105,261) (Fig. 4b, left panel), with more than 72% located in distal regions (Fig. S4b). In contrast, gain of H3K27ac in roots of J411 (clusters C4, C7, and C8, n= 49,527) were much less distributed in distal regions (Fig. 4b, S4b). Of note, many LN-induced H3K27ac-loss loci are specific to J411, but were maintained in KN9204 (C5 and C6 clusters). Such loss of H3K27ac in proximal regions (promoter and genic regions) caused down-regulation of genes that function in auxin homeostasis, cytokinin metabolism, and hormone signaling in J411 under LN/NN conditions (Fig. 4c). However, gain-of-H3K27ac is associated with activation of genes involved in nitrate transport and assimilation, such as *TaNRT2* and *TaNIA* (Fig. 4c). In contrast, gain-of-H3K27ac in roots of KN9204 activated genes involved in cytokinin biosynthesis, auxin polarity transport, transporters for carbohydrate and organic cations, such as *TaD27, TaPILS7*, and *TaLOG4* (Fig. 4c), which function in root growth under LN as reported (Barbez et al., 2012; Tokunaga et al., 2012). Loss of H3K27ac in KN9204 led to down-regulation of genes involved in phosphate ion transport. Consistent with the expression patterns of LN-altered cultivar-specific H3K27ac-associated genes in roots, the root system of J411 has a lower response to LN, whereas LN induced the formation of more root tips and larger root diameters (has a higher LRR-LN response ratio), which eventually increased root surface area and total root volume in KN9204 (Figs. 4d, 4e).

Whether such root morphological change in KN9204 and J411 under LN/NN is regulated by the cultivar-biased LN-induced H3K27ac status? Trichostatin A (TSA), a chemical inhibitor of class I and II histone deacetylase (HDAC) (Yoshida et al., 1990), was used to interfere with the H3K27ac pattern in wheat seedlings grown in a hydroponic culture system (See method for details). The effects of TSA treatment were validated by western blotting (Fig. S4c). In both the TSA treatment and control (untreated), root growth was induced by LN in KN9204, as indicated by increased root tip numbers under LN compared to NN conditions (Fig. 4f). In J411, root growth was not induced by LN under the mock (control) condition (Fig. 4f), as expected. However, root tip numbers increased dramatically by LN in J411 with TSA treatment (Fig. 4f). Thus, TSA treatment inhibited the extensive LN-induced loss of H3K27ac in J411, which in turn restored the root growth response of J411 to LN conditions.

In above-ground tissues, H3K27ac showed more dramatic changes in KN9204 compared to J411, with more peaks gained in flag leaves and more peaks lost in seeds for both KN9204 and J411 under LN/NN conditions (Fig. 4a). In flag leaves, genes with gain-of-H3K27ac were enriched in the sucrose metabolic process and xylem development pathway in KN9204, while genes involved in the response to reactive oxygen species genes were enriched in J411 (Fig. S4d). In seeds, LN mainly induced loss of H3K27ac in both KN9204 and J411 at distal regions instead of gene associated promoters (Fig. S4a). Consistently, reduction in H3K27ac affected few genes in both KN9204 and J411 (Fig. S4e).

Thus, the dynamic changes in H3K27ac that occur in response to low nitrogen conditions preferably enhanced root growth in KN9204 while they strengthened the nitrogen uptake system in J411.

### H3K27me3 shapes distinct root developmental programs in KN9204 and J411 under LN

In addition to H3K27ac, H3K27me3 also showed varied dynamic patterns in roots, flag leaves, and seeds in response to LN in KN9204 and J411(Fig. 5a). In both KN9204 and J411, subtle changes in H3K27me3 occurred in flag leaves, while there were a large number of differential H3K27me3 regions in roots in J411 but in seeds in KN9204 (Fig. 5a). K-means clustering further identified different categories of dynamic H3K27me3 regions in roots and seeds under LN conditions for KN9204 and J411 (Fig. 5b, Fig. S5a).

**Fig. 5.**
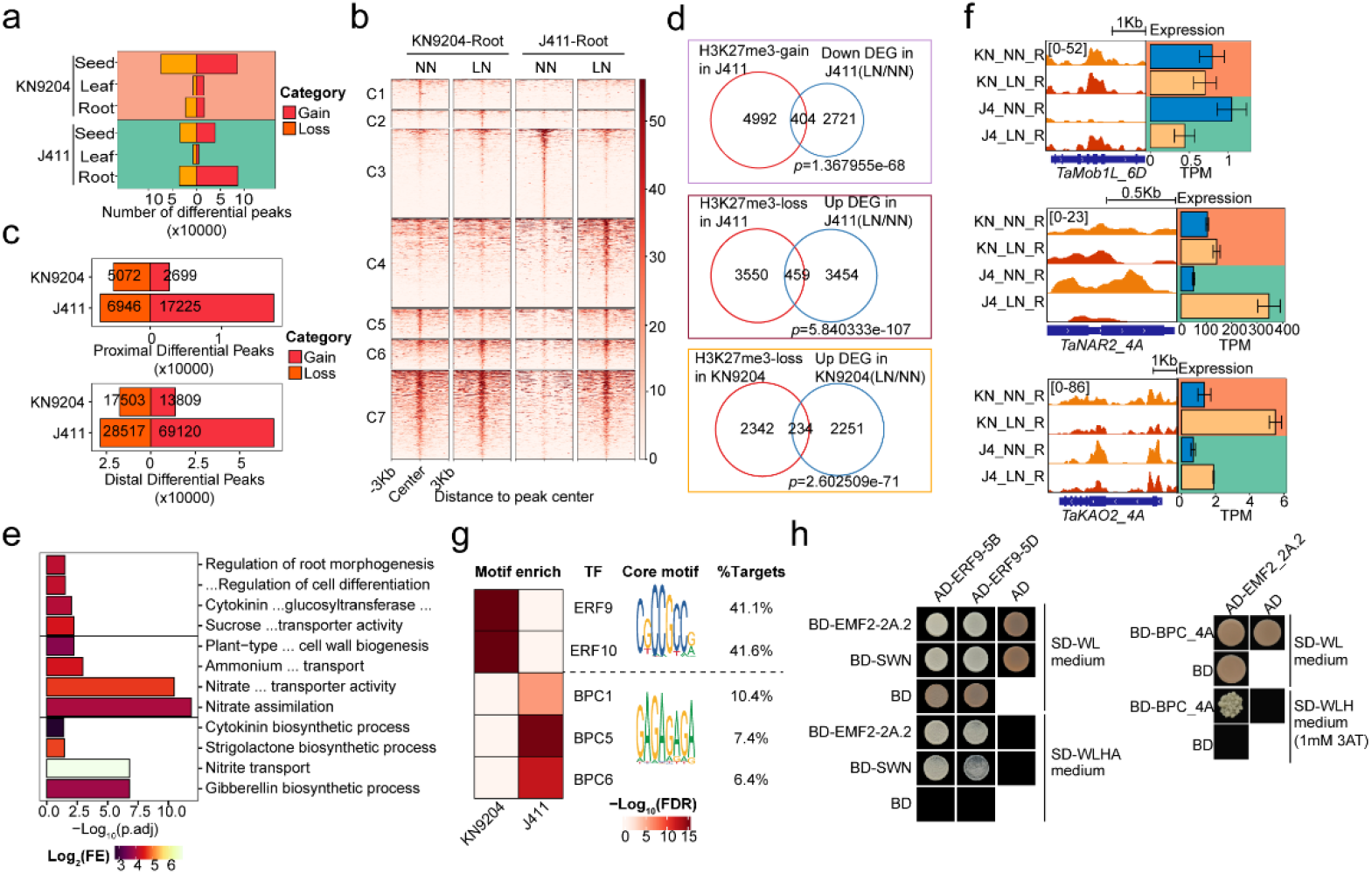
Different trends in H3K27me3 shapes adaptation bias between the cultivars KN9204 and J411. a. The number of differential (gain or loss) H3K27me3 peaks in response to LN in three tissues in KN9204 and J411. b. Heatmaps showing the differential H3K27me3 peaks in the seven clusters in the roots of KN9204 and J411 under LN and NN conditions. c. The number of differential H3K27me3 peaks (in proximal or distal regions) in KN9204 and J411 plants in response to LN. d. Overlap between genes with dynamic changes in H3K27me3 and DEGs in the roots (Fisher’s exact test was used to calculate the *p*-values for the overlaps). e. GO enrichment of the overlapping genes related to (d). f. Representative tracks showing H3K27me3 and transcriptional changes in *TaMob1L_6D* (top panel), *TaNAR2_4A* (middle panel), and *TaKAO2_4A* (bottom panel) for the two cultivars under LN and NN. g. Sequence motif enrichment for the up-regulated H3K27me3 peaks under LN conditions in KN9204 and J411(Fisher’s exact test was used to calculate the *p*-values for the enrichment). h. Yeast two-hybrid (Y2H) assays showing the interaction between TaERF9_5B/TaERF9_5D/TaBPC_4A and TaEMF2-2A.2/TaSWN (components of PRC2). Transformed yeast cells were grown on synthetic media lacking Leu and Trp (−WL) and Leu, Trp, His, and Ade (−WLHA) or Leu, Trp, and His (−WLH) with 1mM 3AT.

The majority of the dynamic H3K27me3 regions were found to be located in distal regions for both roots and seeds (Fig. S5b). In roots, gain-of-H3K27me3 was predominant compared to loss in J411 for both proximal and distal regions (Fig. 5c). There was more loss of H3K27me3 than gain in KN9204 in both proximal and distal regions, although the total peak number was relatively less than in J411 (Fig. 5c). Moreover, genes marked with LN-induced gain-of-H3K27me3 (clusters C4, C5, and C7 in Fig. 5b) significantly overlapped with down-regulated genes in J411 (Fig. 5d, top panel). These genes are enriched in root growth-related processes, including auxin biosynthesis, regulation of cell differentiation, and root morphogenesis (Fig. 5e, top panel). For example, the gene for a *Mob1-like* transcription factor involved in root development (Gonin et al., 2022) has increased H3K27me3 under LN conditions in J411 and the expression level is reduced, while its expression did not change significantly in KN9204 (Fig. 5f, top panel). This indicates that the gain-of-H3K27me3 induced by LN may reduce the root growth response in J411, whereas the gain-of-H3K27me3 in KN9204 only overlapped with 24 genes that were down-regulated by LN in KN9204 (Fig. S5c). Loss-of-H3K27me3 in KN9204 and J411 led to significant overlaps with genes in which expression increased in response to LN (Fig. 5d, middle and bottom panel). Genes involved in the response to nitrite, nitrite transport, and nitrate assimilation were enriched in J411, while genes involved in nitrate transport and SL and GA biosynthetic processes, as well as primary cell wall biogenesis-related genes, were enriched in KN9204 (Fig. 5e, middle and bottom panel). For example, expression of *TaNAR2_4A* and *TaKAO2_4A* was activated separately in J411 and KN9204 with decreased H3K27me3 (Fig. 5f, middle and bottom panel). Thus, LN-induced gain-of-H3K27me3 in roots likely reduces the root growth process, while loss-of-H3K27me3 activates nitrite uptake and metabolism in J411. In contrast, LN-induced H3K27me3 loss in roots tends to activate root growth in KN9204.

Since H3K27me3 deposition in plants is mainly dependent on the recruitment of the “writer” complex Polycomb repressive complex 2 (PRC2) by different DNA recognition factors (Bieluszewski et al., 2021; Xiao et al., 2017), we searched for potential drivers that shape the dynamic H3K27me3 landscape in either KN9204 or J411. Motif scanning analysis found that the CGCCGCC motif (41.1%-41.6%) and the GAGAGA repeat (6.4%-10.4%) were enriched in KN9204- and J411-specific dynamic H3K27me3 regions, respectively (Fig. 5g). Interestingly, those two *cis*-acting motifs were found to be highly enriched in Polycomb response element (PRE) fragments in *Arabidopsis* (Xiao et al., 2017; Zhou et al., 2018). Based on protein-DNA recognition in other species (Fornes et al., 2020), TaERF9_5B/TaERF9_5D and TaBPC_4A were selected for interaction tests with PRC2 components (Fig. 5h). Using yeast two-hybrid (Y2H) and bimolecular fluorescence complementation (BiFC) assays, we verified that TaERF9_9B/TaERF9_5D (AP2/ERF family) and TaBPC_4A could interact with EMF2/SWN (a subunit of PRC2) *in vivo* (Fig. 5h, Fig. S5d). It is worth noting that BPC had been previously reported to interact with SWN and influence root development in *Arabidopsis* (Mu et al., 2017). Thus, LN conditions that induce divergent H3K27me3 changes are likely mediated by different TFs between KN9204 and J411, which contributes to the different developmental routes for the roots of J411 and KN9204 plants.

In the seeds, gain or loss of H3K27me3 in KN9204 did not lead to much change in overall gene expression (Fig. S5d). The same is true for gain-of-H3K27me3 in J411 (Fig. S5e). However, loss-of-H3K27me3 induced a significant amount of up-regulation of gene expression in J411, though no specific GO term was enriched (Fig. S5e).

### Rewiring H3K27me3 modulates root development and nitrate uptake in response to LN

We next asked whether H3K27me3 is functionally important for mediating root development under LN conditions. In total, nine genes encode H3K27me3 methyltransferases within three triads (Fig. S6a), and these genes show varied expression patterns in roots of KN9204 and J411 under different N conditions (Fig. S6b). Considering the high level of expression of *TaSWN* in the roots (Fig. S6b), we used the CRISPR-Cas9 gene editing system to generate knock-out mutants of *TaSWN* (*TraesCS4A02G121300, TraesCS4B02G181400*, and *TraesCS4D02G184600*) in the ‘Bobwhite’ (BW) background (Fig. S6c). Sequencing of transgenic wheat identified a *Taswn-cr* homozygous line with frameshift mutations in all three subgenome copies of *TaSWN* (Fig. S6c). We further used CUT&Tag to profile genome-wide H3K27me3 patterns in the *Taswn-cr* mutant as compared to wild-type BW. The peak number and length of H3K27me3 in *Taswn-cr* was significantly decreased compared to BW (Fig. S6d). Similarly, the intensity of H3K27me3 on coding genes was decreased in *Taswn-cr* compared to BW (Fig. S6e). Therefore, TaSWN is indeed a H3K27me3 writer for certain genomic regions in wheat.

Next, we profiled H3K27me3 patterns in BW and *Taswn-cr* under different nitrogen conditions to identify TaSWN-mediated H3K27me3 deposition in response to LN. K-means clustering identified different categories of dynamic H3K27me3 regions in BW and *Taswn-cr* under NN or LN conditions (Fig. 6a). Of note, the dynamic H3K27me3 peaks were predominantly located in distal regions, especially for clusters 1, 3, 6, and 7 (>80%) (Fig. S6f). Among them, clusters 1 and 6 showed reduced H3K27me3 in *Taswn-cr* compared to BW specifically under LN or both LN and NN conditions (Fig. 6a), which indicated a TaSWN-dependent manner. By overlapping with the LN-induced H3K27me3 peaks in KN9204 and J411 (Fig. 5b), we found that 10%-16% (n=1,748, 14,094 separately) of the LN-induced H3K27me3 peaks in KN9204 or J411 were mediated by TaSWN (Fig. 6b). There were also more genes influenced by SWN in J411 (n=959) compared to KN9204 (n=86) (Fig. 6c, Fig. S6g), suggesting that H3K27me3 regulation has more weight in J411 than in KN9204. Furthermore, we identified a set of genes which gained H3K27me3 in J411 under LN/NN conditions but lost H3K27me3 in *Taswn-cr* compared to BW (Fig. 6d). This group of genes is involved in the root development process; examples are *MADS15, bHLH068*, and *MYB36* (Fig. 6d, Fig. 6e). The expression of these genes was induced in *Taswn-cr* under LN/NN, but there were no changes in BW (Fig. 6f). These results hint that TaSWN-mediated LN-induced gain-of-H3K27me3 in J411 partially reduces the root growth response to LN conditions. Therefore, we speculated that root growth in *Taswn-cr* plants would be more responsive to LN compared to BW. Indeed, we found roots were more developed in *Taswn-cr* compared to BW plants under LN, as determined by total root length and the number of root tips (Fig. 6g, Fig. 6h). In addition, the relative level of induction of *NRT2* expression in response to LN in *Taswn-cr* plants was lower compared to BW (Fig. 6i, left panel), which were similar to KN9204 plants with more developed root systems (Fig. 6i, right panel).

**Fig. 6.**
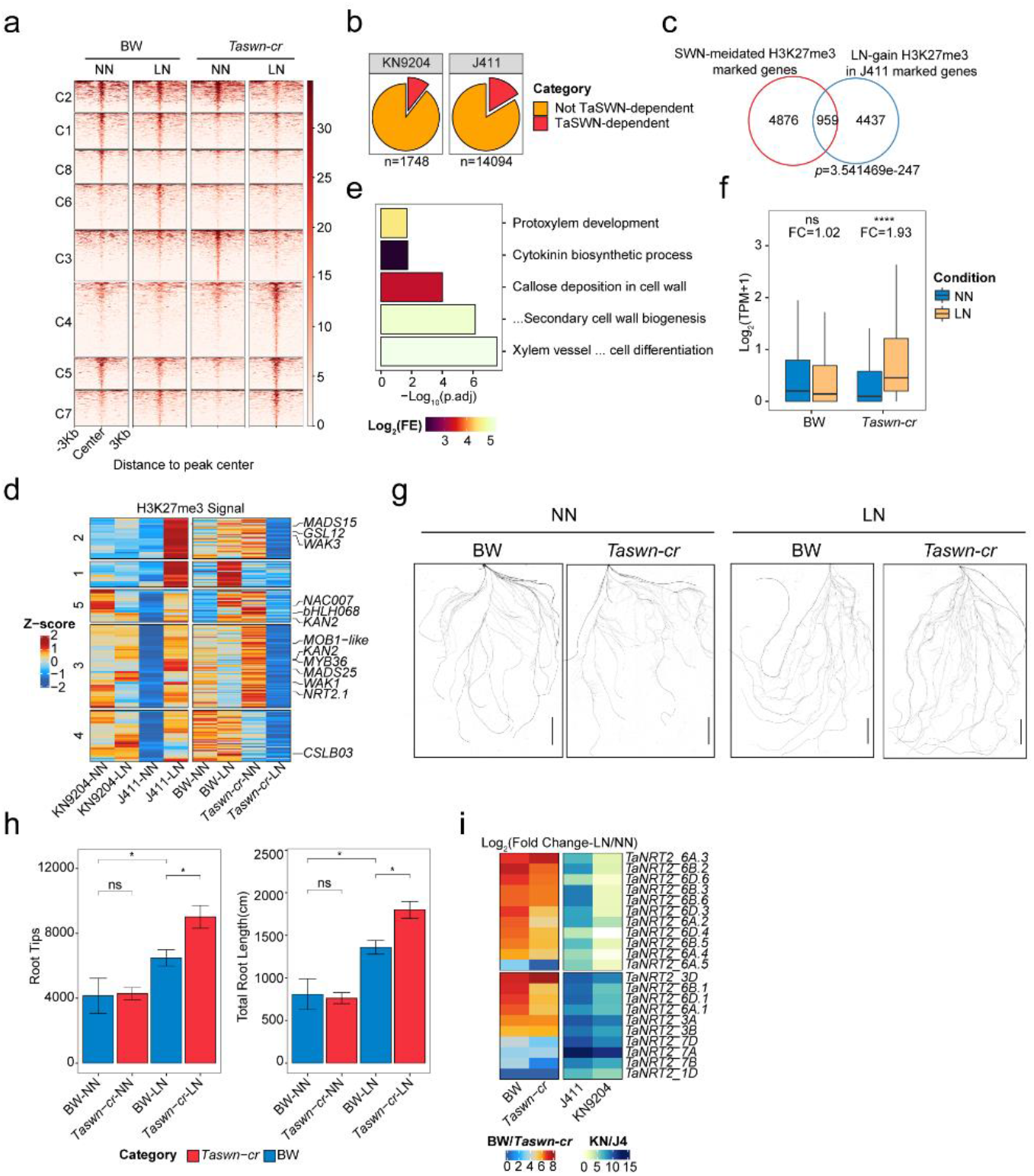
Rewiring H3K27me3 strengthens root adaptation to low nitrogen fertilization level. a. Heatmaps showing the differential H3K27me3 peaks in ‘BobWhite’ (BW) and the *Taswn-cr* mutant plants in the eight clusters under NN and LN conditions. b. The proportion of TaSWN-dependent peaks in the up-regulated H3K27me3 peaks under LN conditions in KN9204 and J411. c. Overlap between genes marked by TaSWN-dependent H3K27me3 and marked by gain-of-H3K27me3 under LN conditions in J411 (Fisher’s exact test was used to calculate the *p*-values for the overlap). d. Dynamic changes in the TaSWN-dependent H3K27me3 peaks under different nitrogen conditions (NN and LN) in KN9204, J411, BW, and *Taswn-cr* plants. Representative gene names are shown on the right. e. GO enrichment of the overlapping genes related to (d). f. The expression changes and fold-changes of the genes in (d). (Wilcoxon test: ns: *p*>0.05; ****: *p*≤0.0001) FC: fold-change. g. Scans of root systems of BW and *Taswn-cr* seedlings grown under different nitrogen conditions. Scale bars=5 cm. h. The number of root tips and total length of the root systems for BW and *Taswn-cr* seedlings grown under different nitrogen conditions. (Student’s *t*-test; ns: *p*>0.05; *: *p*≤0.05). i. The fold-changes in the expression of *NRT2* genes in BW, *Taswn-cr*, J411, and KN9204 seedlings grown under different nitrogen conditions.

The comparison of root morphological changes and transcriptional profiles in response to LN between BW and the *Taswn-cr* mutant highlighted the fact that H3K27me3 plays important role in balancing root growth and nitrogen metabolism in response to LN conditions. Rewiring H3K27me3 could affect the plant strategy in selecting between developing root growth and raising the nitrogen uptake system to coordinately adapt to low nitrogen conditions.

## Discussion

As the urgency to reduce nitrogen fertilizer application in crop production, considerable effort has been directed towards dissecting the genetic basis of NUE regulation in crops (Han et al., 2015; Li et al., 2017; Swarbreck et al., 2019). However, epigenetic regulation, which functions in coordinating with transcription factors to manipulate gene expression, is not well studied in wheat. To fill this gap, we generated epigenomic datasets for three different tissues in two wheat cultivars that differ with respect to NUE (KN9204 and J411) under different nitrogen conditions (Fig. 1). Our analysis revealed that the epigenome, which varies more than DNA sequence variation, plays an important role in mediating the cultivar-specific low nitrogen response.

### Epigenetic variation contributes to cultivar-specific trait formation

Epigenetic modifications, especially H3K27me and H3K27ac, mediate transcriptional dynamics and contribute to different developmental programs or nitrogen metabolic processes between cultivars (Figs. 1, 2). Bias-expressed NMGs are associated with altered epigenetic regulation patterns rather than DNA sequence variations between KN9204 and J411 (Fig. 2). In addition, distal regulatory regions (H3K27ac and H3K27me3) clearly reflect cultivar specificity, with higher DNA sequence variations, and are associated with previously identified NUE-related QTLs (Fig. 3). In maize, many distal regulatory regions have been reported to regulate gene expression and are associated with agronomic trait variations (Li et al., 2019; Ricci et al., 2019). Indeed, genes associated with cultivar-specific H3K27ac either in the promoter or in the distal region are enriched in QTL regions for mediating NUE-related agronomic traits, such as MRL, GPC, and Nup (Fig. 3), which could be good candidates for mediating the genetic difference between KN9204 and J411. Regarding the important role of epigenetic regulatory regions, more cultivars with defined NUE features could be used to profile the epigenome, especially H3K27ac and H3K27me3 in the future, which would enable Epi-GWAS analysis to uncover NUE regulatory mechanisms.

### Epigenetic regulation balances root growth and nitrogen metabolism under LN constraint

To absorb adequate nitrogen in nitrogen-limiting conditions, cultivars with different NUE features have various strategies, such as triggering root growth to have more root tips and increase the total root system volume in KN9204, or powering up the nitrate uptake machinery via up-regulation of NRT2 transporters in J411 (Fig. 1). Interestingly, the different adaptive strategies in roots between KN9204 and J411 are correlated with changes in the dynamic epigenome, especially for H3K27ac and H3K27me3 (Figs. 4, 5). Gain-of-H3K27ac and loss-of-H3K27me3 coordinately enhance the expression of root development-related genes in KN9204 under LN conditions, whereas loss-of-H3K27ac and gain-of-H3K27me3 reduce root development in J411 under LN, but rather activate nitrate uptake transporters via gain-of-H3K27ac and loss-of-H3K27me3 (Fig. 7). Several transcription factors show the potential to establish such epigenetic modification specificity via the recognition of certain cis-acting motifs and recruiting histone modification writers or erasers, such as ERFs and BPCs (Fig. 5). Thus, precise epigenetic modification modulates root development and nutrient absorption, which is likely to be a general mechanism for balancing plant growth and environmental stimulus response or adaptation (Xiao et al., 2022). However, it is presently unknown how such epigenetic modification behaves in a cultivar-specific manner. An interesting finding is that the potential histone writer or eraser-guiding TFs are located within the QTL regions linked to the MRL and Nup traits (Supplemental Table 3). Further analysis would elucidate how LN can trigger different response patterns in those TF genes in cultivars that vary with respect to NUE.

**Fig. 7.**
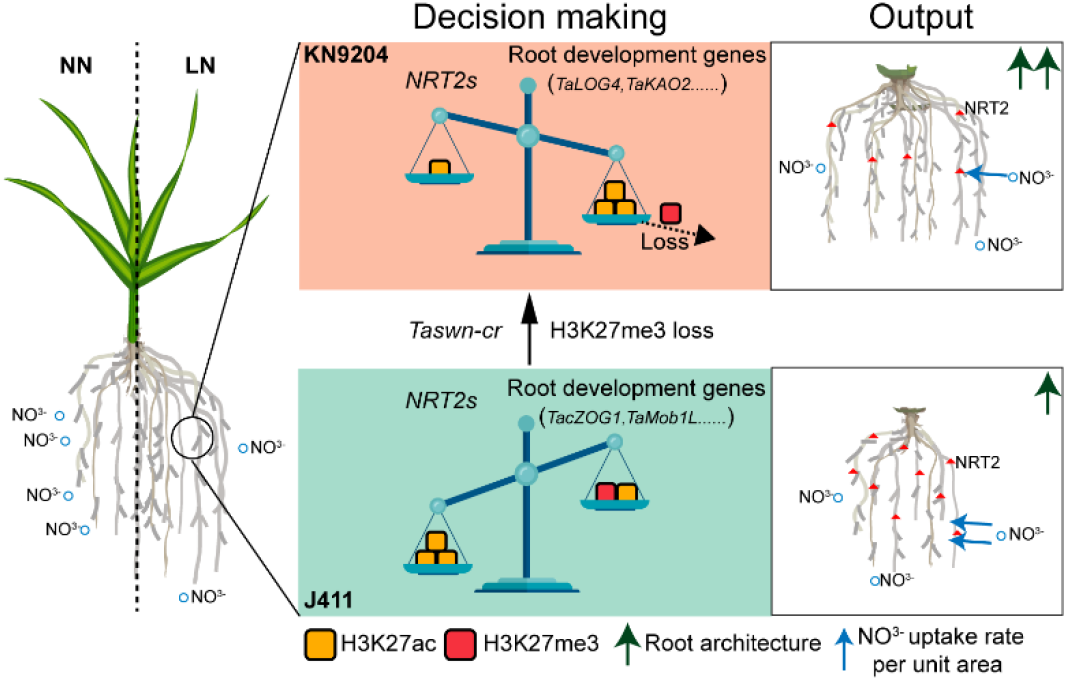
The balanced model of epigenetic regulation for divergent strategies to LN conditions in wheat.

### Manipulating epigenetic regulation to decouple root growth and nitrogen metabolism for NUE improvement in wheat

In addition to the correlation between cultivar-specific epigenetic dynamics and varied strategies in response to LN, we have shown that manipulating the epigenetic features could affect the strategy selection for LN adaptation. Chemical inhibition (TSA treatment) or genetic manipulation (*Taswn-cr*) that changes the epigenome landscape (H3K27ac and H3K27me3) could lead to altered root system development and coordinated *NRT2* induction intensity under LN constraint (Figs. 4, 6). Similarly, several reports have shown that adjusting histone modification contributes to the response and/or tolerance to stress, such as HDA6-regulated salt stress (Luo et al., 2012), and JMJ1-regulated dehydration stress (Huang et al., 2019). Interestingly, the enhancement of root growth is likely coupled with the attenuated induction of the expression of nitrate transporter coding genes under LN constraint by histone modification, in particular H3K27me3 (Fig. 7). Theoretically speaking, precise epigenomic modification alterations at specific regions/genes could help to decouple such linkage, which could generate wheat with developed root architecture system and higher induction of nitrogen transporters simultaneously. To achieve this, instead of the genetic manipulation of the “writer” or “eraser” to histone modification, fine-tuning of the driver (for example ERFs) may serve as a better way of coordinating nitrogen metabolism and adaptive root growth to low nitrogen constraints in wheat cultivars.

## Materials and methods

### Plant materials and culture conditions

The root of KN9204 and J411 was harvested 4 weeks after transplanting in the nutrient solution, which corresponding to 28-day in previous study (Shi et al., 2022), and immediately frozen in liquid nitrogen and stored at -80°C. The nutrient solution for NN was as follows: 1 mM Ca(NO_3_)_2_, 0.2 mM KH_2_PO_4_, 0.5 mM MgSO_4_, 1.5 mM KCl, 1.5 mM CaCl_2_, 1 × 10^−3^ mM H_3_BO_3_, 5 × 10^−5^ mM (NH_4_)_6_Mo_7_O_24_, 5 × 10^−4^ mM CuSO_4_, 1 × 10^−3^ mM ZnSO_4_, 1 × 10^−3^ mMMnSO_4_, 0.1 mM Fe(III)–EDTA. For LN, 0.02mM Ca(NO_3_)_2_, 2.48 mM CaCl_2_ (to compensate for the Ca^2+^ concentration in the nutrient solution), and other component was not changed. The flag leaf was harvested at heading stage (Shi et al., 2022) in the field (Shijiazhuang, China), and seed was also harvested 21DAA (Shi et al., 2022) in the field (shijiazhuang, China). In each NN plot, 300 kg/ha of diamine phosphate and 225 kg/ha of urea were applied before sowing, and 150 kg/ha of urea was applied at the elongation stage every year. In the LN plots, no N fertilizer (N-deficient) was applied during the growing period.

### Generation of transgenic wheat plants

To obtain CRISPR transgenic wheat plants, the *pU6-gRNA* of *TaSWN* was annealed and inserted into *pJIT163-Ubi-Cas9* vector. All constructed vectors were transformed into callus to generate the transgenic plants.

To identify mutations in *TaSWN-4A, TaSWN-4B*, or *TaSWN-4D*, gene-specific primers were designed around the target site. Primers F and A-R were used to amplify *TaSWN-4A*, F and B-R were used to amplify *TaSWN-4B*, and F and D-R were used to amplify *TaSWN-4D*. PCR products were checked on agarose gels and genotyped by Sanger sequencing.

### RNA-seq and CUT&Tag experiment

Total RNA was extracted using HiPure Plant RNA Mini Kit according to the manufacturer’s instructions (Magen, R4111-02), and libraries were sequenced using an Illumina Novaseq platform.

CUT&Tag experiment were done follow the previous described method (Zhao et al., 2023). The nuclei were extracted by chooping fresh samples soaked in the HBM buffer (25 mM Tris-HCl pH 7.6, 0.44 M sucrose, 10 mM MgCl2, 0.1% Triton-X, 10 mM Beta-mercaptoethanol, 2 mM spermine, 1 mM PMSF, EDTA-free protease inhibitor cocktail). After overnight incubation with corresponding antibody in 4°C, the nuclei was incubated in 50μl wash buffer (20 mM HEPES pH 7.5; 150 mM NaCl; 0.5 mM Spermidine; 1× Protease inhibitor cocktail) with secondary antibody (1:100; Guinea Pig anti-Rabbit IgG antibody) at 4°C for around 1-2 hour and then washed twice with wash buffer. pA-Tn5 complex (A 1:100 dilution in CT-300 buffer:20 mM HEPES pH 7.5; 300 mM NaCl; 0.5 mM Spermidine; 1× Protease inhibitor cocktail) was incubated with nuclei in 4°C for 2-3h. After washing twice with CT-300 buffer, the tagmentation of nuclei was done in 300 μl Tagmentation buffer (20 mM HEPES pH 7.5; 300 mM NaCl; 0.5 mM Spermidine; 1× Protease inhibitor cocktail; 10 mM MgCl2) in 37°C for 1h. 10 μl 0.5M EDTA, 3 μl 10% SDS and 2.5 μl 20 mg/ml Protease K were added to stop tagmentation reaction. The DNA was extracted with phenol:chloroform:isoamyl alcohol, precipitated with ethanol, and resuspended in ddH2O. The library was amplified 17 cycles by Q5 high fidelity polymerase (NEB, M0491L). Antibodies used for histone modifications are the same as previous reported (Zhao et al., 2022a). Libraries were purified with AMPure beads (Beckman, A63881) and sequenced using the Illumina Novaseq platform at Annoroad Gene Technology.

### Nitrate assays

Nitrate assays were performed as described (Zhao and Wang, 2017). Standard curve was made based on different concentration of KNO_3_ solution (deionized water as a control). Samples were boiled at 100 °C for 20 min. After Centrifuge of the boiled different samples, salicylic acid-sulphuric acid was added to supernatant. After incubation of 20 min, 8% (w/v) NaOH solution was added. After cool down of the samples, measure the OD410 value of each sample with the control for reference.

For samples collected of KN9204 and J411, grind each sample into powder in liquid nitrogen, detection procedure was done as described before. Finally, calculate the nitrate concentration using the following equation: Y = CV/W.

Y: nitrate content (μg/g),

C: nitrate concentration calculated with OD410 into standard curve (μg/ml),

V: the total volume of extracted sample (ml),

W: weight of sample (g).

### Grain protein content assays

Grain protein content was measured by near-infrared reflectance spectroscopy (NIRS) with a Perten DA-7200 instrument (Perten Instruments, Huddinge, Sweden) and expressed on a 14 % moisture basis. The measurements were calibrated using calibration samples according to the manufacturer’s instructions.

### Root system scanning

The root of different samples was scanned by ScanMaker i8000 plus, after analyzed by WinRHIZO software, five root traits were quantified, including total root length (Rl), root surface area (Rs), root volume (Rv), root diameter (Rd) and root tip number (Rt); and maximum root length is measured using a ruler.

LN response ratio (LRR) was calculated to reflect the root change under LN condition, which was (R_LN_(root trait under LN condition) - R_NN_(root trait under NN condition))/R_NN_.

### TSA treatment

TSA (V900931-5MG) was dissolved in DMSO, then directly add into nutrient solution (NN/LN) to final concentration of 2 μM, with DMSO as mock. After treatment of 4 days, the root was harvested and immediately frozen in liquid nitrogen and stored at -80°C.

### Protein interaction test (Y2H + BiFC)

For the interaction of TaHD2_1D/TaHAG2_5A and TaDREB26_6B, Full-length of *TaDREB26_6B* were amplified using specific primers and fused with GAL4 AD in the *pDEST22* vector. Full length of *TaHD2_1D* and *TaHAG2_5A* were amplified using specific primers and fused with GAL4 BD in the *pDEST32* vector. Interactions in yeast were tested on the SD/-Trp/-Leu/-His/ medium with 2.5 mM 3AT.

For the interaction between PRC2 and TaERF9_5B/ TaERF9_5D, Full-length of *TaERF9_5B* and *TaERF9_5D* were amplified using specific primers and fused with GAL4 AD in the *pDEST22* vector. Full length of *TaEMF2-2A.2* and N-terminal of *TaSWN* were amplified using specific primers and fused with GAL4 BD in the *pDEST32* vector. Interactions in yeast were tested on the SD/-Trp/-Leu/-His/-Ade medium.

For the BiFC analysis, the cDNA of *TaERF9_5B/ TaERF9_5D/TaBPC_4A*and *TaEMF2-2A.2/TaSWN* was amplified with primers and cloned into *pSCYCE* and *pSCYNE* vectors containing either C- or N-terminal portions of the enhanced cyan fluorescent protein. The resulting constructs were transformed into *A. tumefaciens* strain GV3101. Then these strains were injected into tobacco leaves in different combinations with p19. The CFP fluorescence was observed with a confocal laser-scanning microscope (FluoView 1000, Olympus).

### Western blot assays

Total histone proteins were extracted by using EpiQuik Total Histone Extraction Kit (OP-0006-100). The total histone proteins were then used for western blot using the antibodies listed below. Anti-H3 immunoblot was used as a loading control. Antibodies: anti-H3 (ab1791, Abcam), anti-H3K27ac (ab4729, Abcam). Immunoblotting was done by using the enhanced chemiluminescence (ECL) system.

### RNA-seq Data Processing

Adapter sequence and low-quality reads of RNA-seq library was removed by fastp (0.20.1)(Chen et al., 2018), the cleaned reads was mapped to IWGSC Refseq v1.1 using hisat2 (2.1.0)(Kim et al., 2019), and gene expression was quantified by featureCount (2.0.1)(Liao et al., 2014). Differentially expressed genes were evaluated using the DESeq2 package (1.34.0)(Love et al., 2014) in R with an adjusted p value < 0.05 and log2 fold-change > 1. TPM (Transcripts Per Kilobase Million) values generated from the counts matrix were used to characterize gene expression and used for hierarchical clustering analysis.

For functional enrichment, GO annotation files were generated from IWGSC Annotation v1.1 and an R package clusterProfiler (4.2.2)(Wu et al., 2021) was used for enrichment analysis.

### CUT&Tag Data Processing

Adapter sequence and low-quality reads of CUT&Tag library was removed by fastp (0.20.1)(Chen et al., 2018), the cleaned reads was mapped to IWGSC Refseq v1.1 using bwa mem algorithm (0.7.17)(Li, 2013), We further filter the reads mapped with “samtools view -bS -F 1,804 -f 2 -q 30” to filter the low-quality mapped reads. Then the high-quality mapped reads were reduplicated using Picard-2.20.5-0. The de-duplicated bam files from two biological replicates were merged by samtools (1.5)(Danecek et al., 2021), and merged bam file was converted into bigwig files using bamCoverage provided by deeptools (3.3.0) with parameters “-bs 10 --effectiveGenomeSize 14,600,000,000 -- normalizeUsing RPKM --smoothLength 50”. The bigwig files were visualized using deeptools (3.3.0)(Ramirez et al., 2016) and IGV (2.8.0.01)(Robinson et al., 2011).

For peak calling, macs2 (2.1.4)(Zhang et al., 2008) was used. For narrow peaks (H3K27ac, H3K4me3, and H2A.Z) and broad peaks (H3K27me3, H3K36me3, and H3K9me3), parameters “-p 1e-3 --keep-dup all -g 14600000000” and “--keep-dup all -g 14600000000 --broad --broad-cutoff 0.05” were used. Peak was annotated to the wheat genome using the R package ChIPseeker (v1.30.3)(Yu et al., 2015), as peaks annotated to three categories: promoter (−3000bp of TSS), genic (TSS to TES) and distal (other). The MAnorm package(Shao et al., 2012) was used for the quantitative comparison of CUT&Tag signals between samples with the following criteria: |M value| > 1 and P < 0.05.

### Chromatin state analysis

For chromatin state analysis, chromHMM (1.21)(Ernst and Kellis, 2017) was used. “BinarizeBam” and “LearnModel” commands with default parameters were used for chromatin-state (CS) annotation. Multiple models were trained on these data, with CS numbers ranging from 2 to 20. The 15-state model was selected because it captured all the key information of CS. In previous studies, 15-state models were similarly trained on rice (Zhao et al., 2020) and *Arabidopsis* (Wu et al., 2022) data.

For chromatin states dynamic change analysis, bins (CS called, 200bp) were called dynamic if its state diverges between different samples. For variability score of histone modification, one minus jaccard index, which was calculated by bedtools (v2.29.2).

### Distal regulatory region-gene assignment

The distal regulatory regions annotation strategy was largely based on a previous study (Trevino et al., 2020). Genes within 0.5M from a distal H3K27ac peak are considered candidate target genes. Then we generated null model as correlations between randomly selected peaks and randomly selected genes on different chromosomes, and enabling us to compute mean and standard deviation of this null distribution. For each potential link, after calculate correlation between gene expression (TPM) and distal H3K27ac signal (FPKM) in samples, we also compute p-values for the test correlations based on null model, then significantly pairs were selected as regulatory region-gene pairs.

### Detection of transcription factor-binding motifs

To detect recruiter of dynamic H3K27me3 changes, we downloaded the position weightmatrices of plant motifs from the JASPAR database(Fornes et al., 2020), the motifs was scanned by FIMO(4.11.2)within dynamic H3K27me3 regions. And enrichment test of motifs detected were done by fisher test in R.

### Statistical analysis

R (https://cran.r-project.org/;version 4.2.1) was used to compute statistics and generate plots if not specified. For two groups’ comparison of data, the student’s t-test was used, such as Fig.2g, Fig.2i, Fig.4d, Fig.4e, Fig.4f, Fig.6h. For enrichment analysis, Fisher’s exact test was used, such as Fig.2b, Fig.3c, Fig.3f, Fig.5d, Fig.5g, Fig.6c, Fig.S3b, Fig.S4e, Fig.S5e, Fig.S6g.

## Data availability

The raw sequence data were deposited in the Genome Sequence Archive (https://bigd.big.ac.cn/gsa) under accession number PRJCA014417; the transcriptome used was under BioProject accession numbers PRJCA004416; Hi-C data of wheat root was download from Gene Expression Omnibus (GEO) under accession number GSM3929164.

## Acknowledgements

This research was supported by National Key Research and Development Program of China (2021YFF1000401, 2021YFD1201500), National Natural Sciences Foundation of China (U22A6009), the Strategic Priority Research Program of the Chinese Academy of Sciences (XDA24010104) and China Agriculture Research System of MOF and MARA (CARS-03).

## Author contributions

J.X. designed and supervised the research, J.X. H.Z. J.-M. L. H.-Q.L. wrote the manuscript. H.Z. performed CUT&Tag, RNA-seq, nitrate assay, TSA treatment, yeast two-hybrid assay and root scanning experiments; H.Z. and L.Z. performed data analysis. X.-Y.Z. performed plasmid construction, part of yeast two-hybrid assay, BiFC experiment. F.C. performed determination of grain protein content. J.Z. performed Western blot; J.-C.C. performed plasmid construction for *Taswn-cr*. C.-X.G. provided *Taswn-cr* transgenic wheat. H.Z. and J.X. prepared all the figures. L.W., X.-D.F., and Y.-P.T. polished the manuscript. All authors discussed the results and commented on the manuscript.

---

**Fig. S1.**
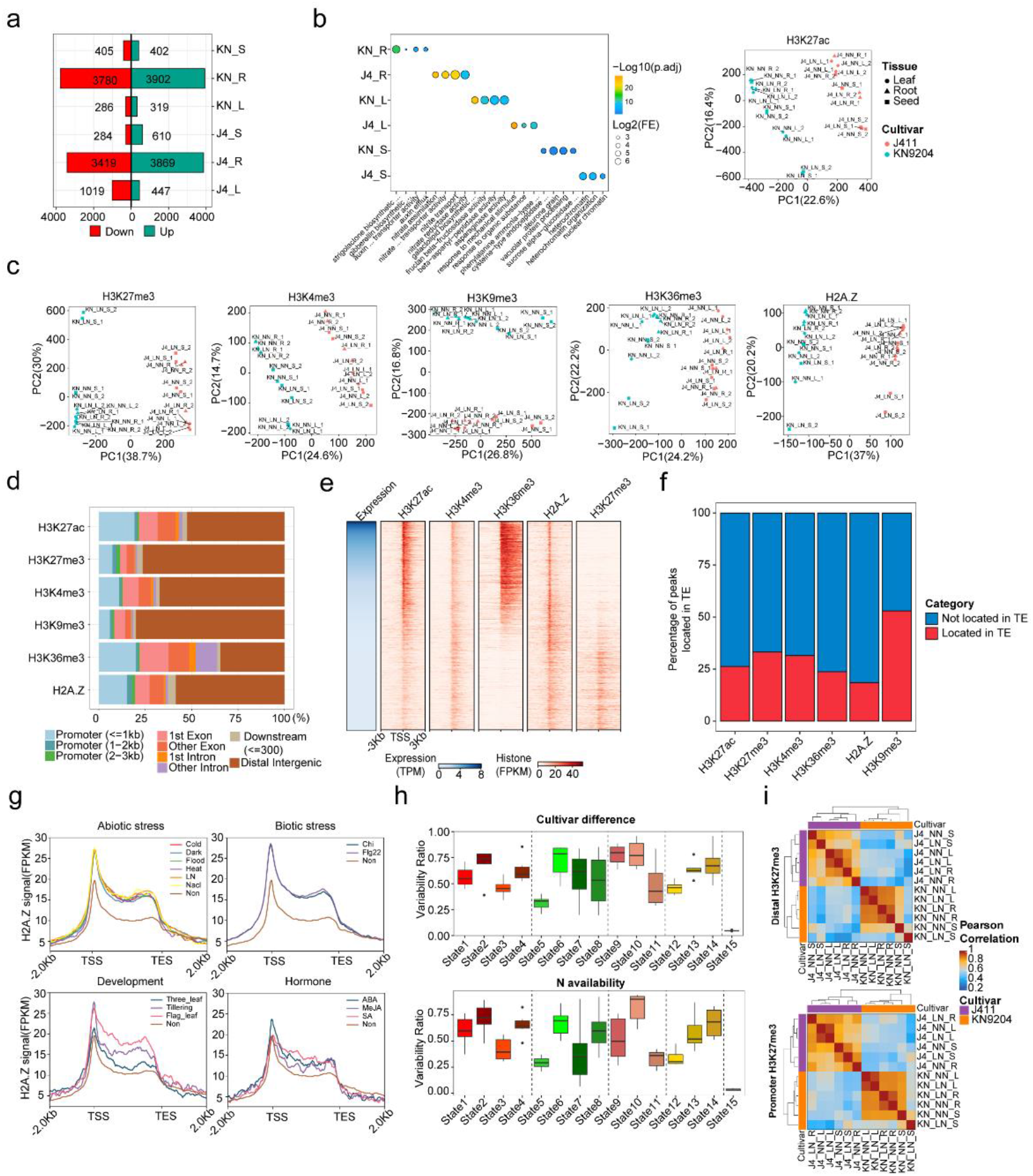
NUE epigenome dataset. a. The number of DEGs in the tissues in response to LN in KN9204 and J411. b. GO enrichment of cultivar-specific LN-induced DEGs. c. PCA plots of H3K27ac, H3K27me3, H3K4me3, H3K9me3, H3K36me3, and H2A.Z in the NUE epigenome dataset. Each dot represents one sample. Two biological replicates were sequenced for each stage. d. Peak distributions in gene regions of different histone marks in the wheat genome. e. Heatmaps of epigenetic marks for all annotated wheat genes which were sorted according to their expression levels determined by RNA-Seq. f. Proportion of peaks located in TE regions for the six different histone marks. g. H2A.Z profiles of genes that respond to external stimuli (abiotic and biotic stress); RNA-seq data from (Wang et al., 2021). h. Fraction of bases for 15 states that vary between the different cultivars (top) and nitrogen availability (bottom). i. Cross-correlation heatmaps of all H3K27me3 peaks which are located in distal or promoter regions separately.

**Fig. S2.**
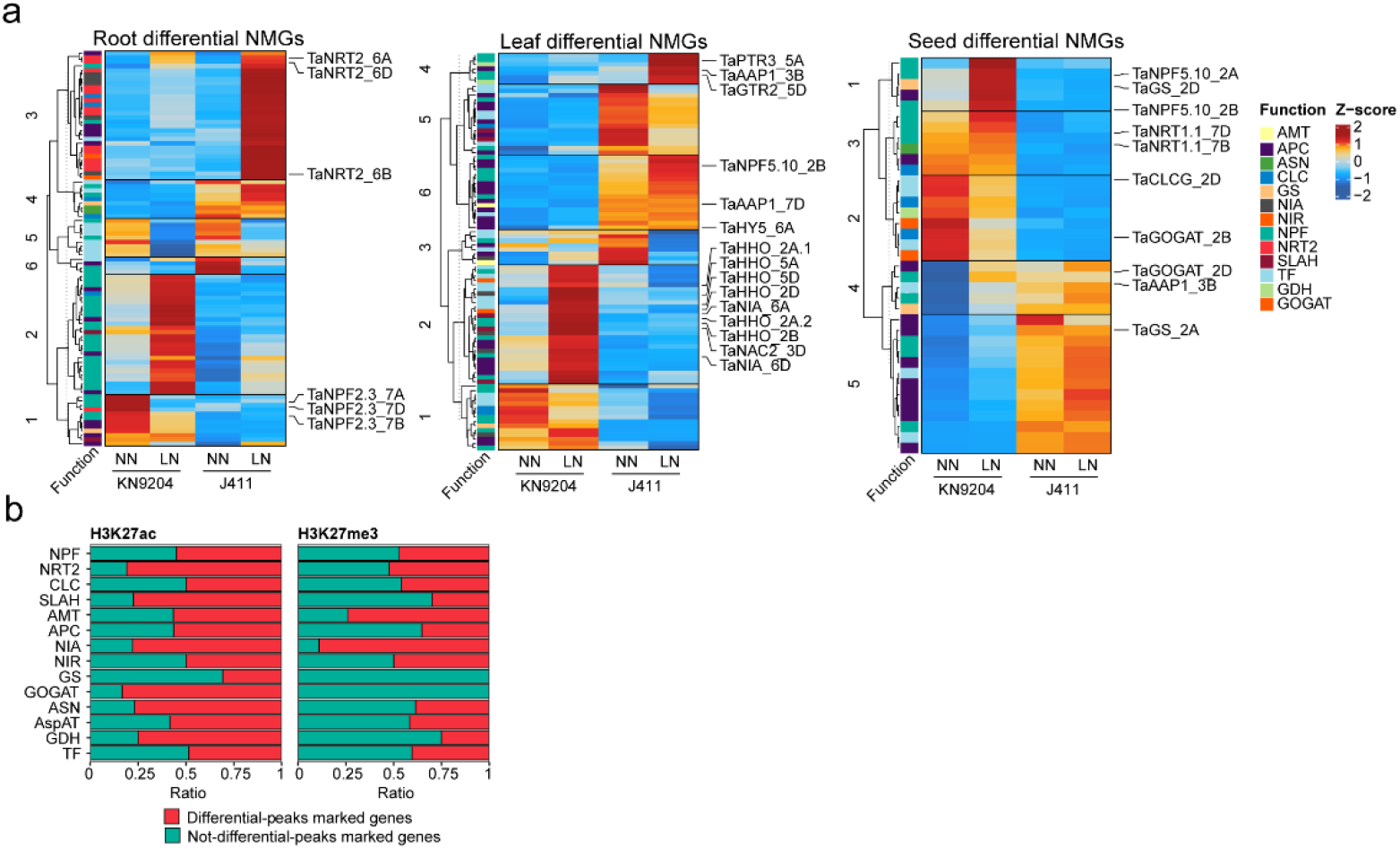
Dynamic changes in the transcriptomes and epigenomes of NMGs. a. K-means clustering of LN-induced differentially-expressed NMGs in roots, leaves, and seeds. See also Supplemental Figure S1, Dataset I showing the list of LN-induced differentially-expressed NMGs in the three tissues, Dataset II showing the list of NMGs marked by differential H3K27ac, and Dataset III showing the list of NMGs marked by differential H3K27me3. b. Percentage of NMGs in different categories marked by differential H3K27ac and H3K27me3.

**Fig. S3.**
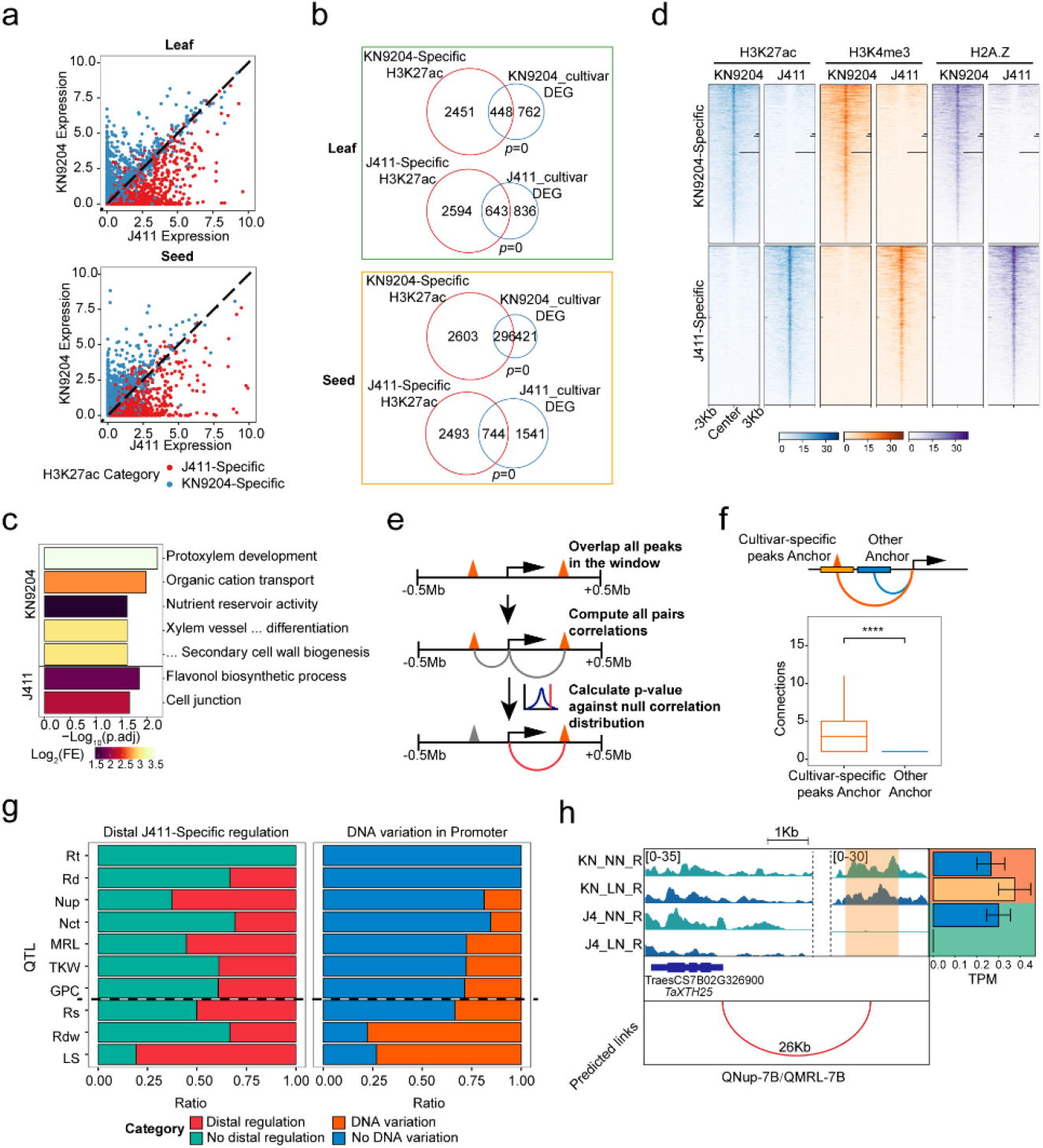
The functional influence of cultivar-biased H3K27ac peaks. a. Mean gene expression in J411 (x-axis) versus mean gene expression in KN9204 (y-axis) for genes associated with proximal cultivar-specific H3K27ac peaks separately in the leaf and seed. b. Overlap between genes marked by cultivar-specific H3K27ac and DEGs between cultivars (blue circle) in the leaf (top panel) and seed (bottom panel) (Fisher’s exact test was used to calculate the *p*-values for the overlaps). c. GO enrichment of genes that are marked by proximal cultivar-specific H3K27ac peaks in KN9204 and J411. d. Heatmaps showing the co-localization between cultivar-specific H3K27ac peaks and H3K4me3/H2A.Z. e. Schematic diagram showing the approach used to link distal cultivar-specific H3K27ac peaks to genes. f. Cross validation of distal cultivar-specific H3K27ac peaks assigned by Hi-C data. g. Fraction of DEGs with distal J411-specific regulation or DNA variation within the promoter located in QTLs between KN9204 and J411. Abbreviation are as given in Fig. 3e. h. Representative tracks of *TaXTH25* regulated by distal cultivar-specific H3K27ac peaks in *QMRL-7B*.

**Fig. S4.**
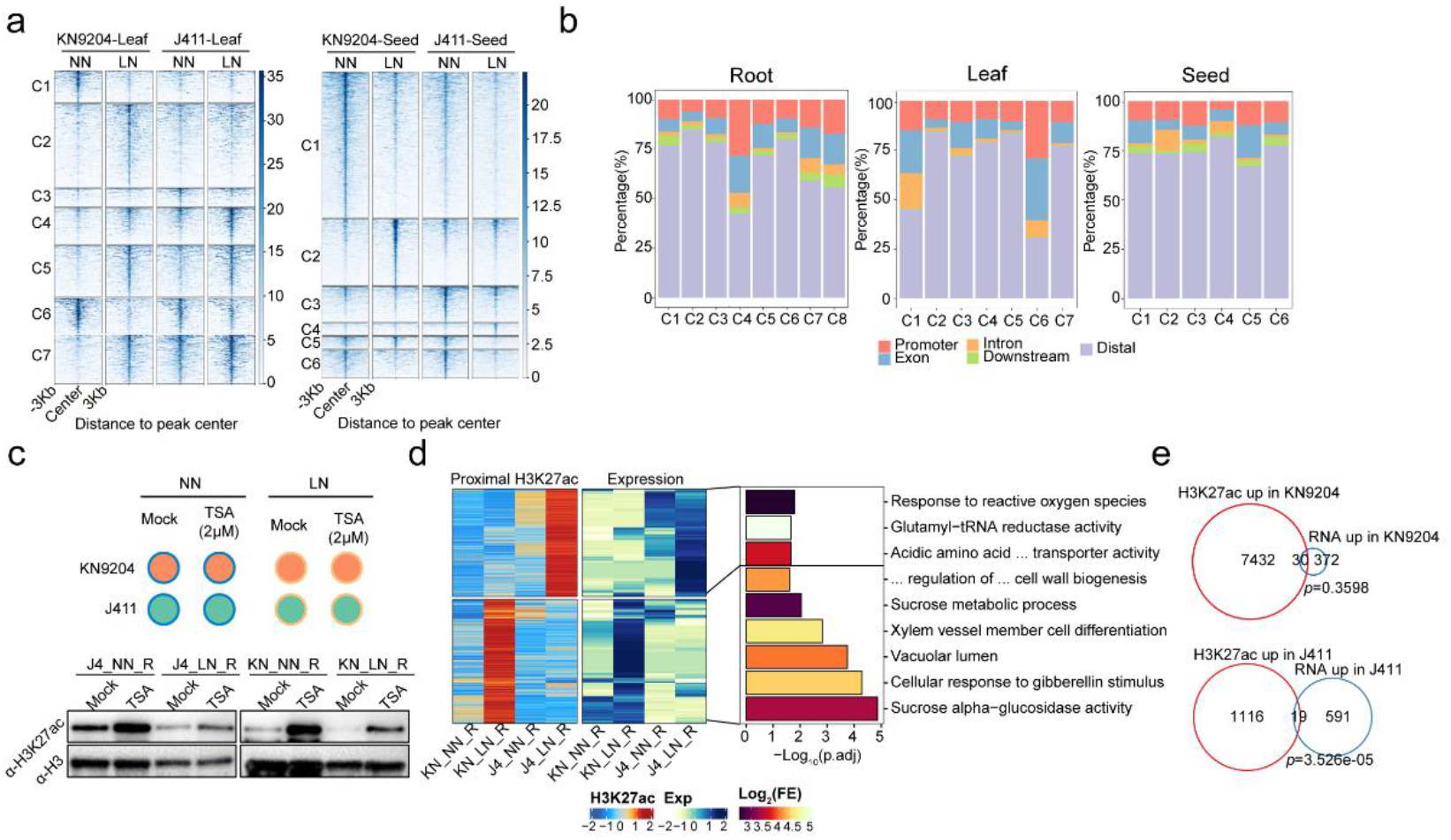
Dynamic H3K27ac changes in KN9204 and J411. a. Heatmaps showing the differential H3K27ac peaks in the leaves and seeds of KN9204 and J411 under LN and NN conditions. b. Peak distribution of differential H3K27ac peaks in roots, leaves, and seeds. c. Experiment design and western blotting of TSA (2 μM) treatment of KN9204 and J411 seedlings under the two nitrogen conditions. d. Dynamic H3K27ac and corresponding expression changes in the leaves of KN9204 and J411 seedlings. GO enrichments are shown on the right. e. Overlap between genes with up-regulated H3K27ac and the up-regulated DEGs for KN9204 and J411 in the seeds (Fisher’s exact test was used to the calculate *p*-values for the overlaps).

**Fig. S5.**
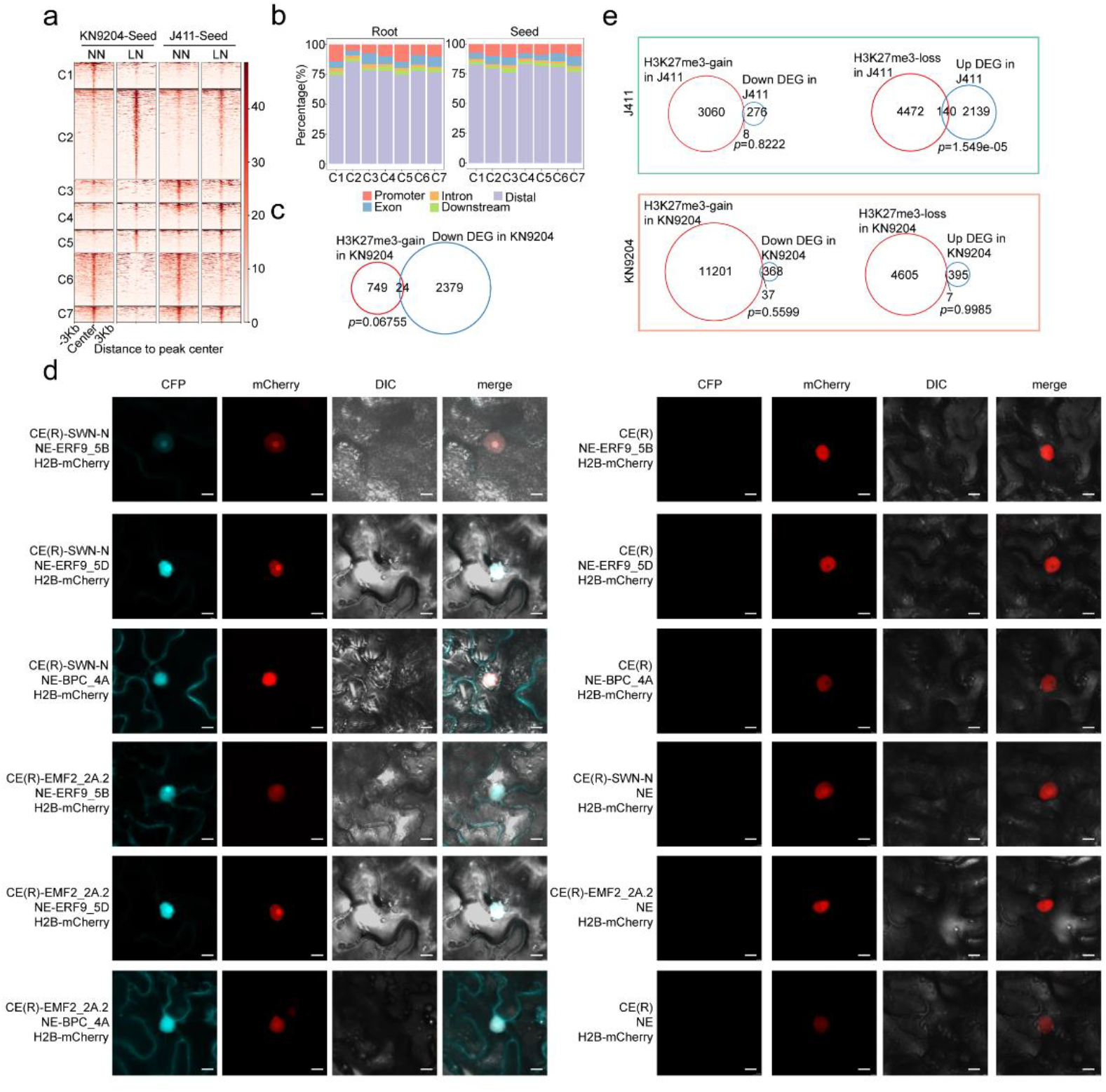
Dynamic H3K27me3 changes in KN9204 and J411. a. Heatmaps showing the differential H3K27me3 peaks in seven clusters in the seeds of KN9204 and J411. b. Distribution of differential H3K27me3 peaks in the roots and seeds. c. Overlap between genes up-regulated for H3K27me3 and the down-regulated DEGs in the roots of KN9204 (Fisher’s exact test was used to calculate the *p*-values for the overlaps). d. BiFC analysis of the interaction between TaERF9_5B/TaERF9_5D/TaBPC_4A and TaSWN-N/TaEMF2_2A.2. Scale bars, 10 mm. e. Overlap between genes up-regulated for H3K27me3 and the down-regulated DEGs in the seeds of J411 (top panel) and KN9204 (bottom panel) (Fisher’s exact test was used to calculate the *p*-values for the overlaps).

**Fig. S6.**
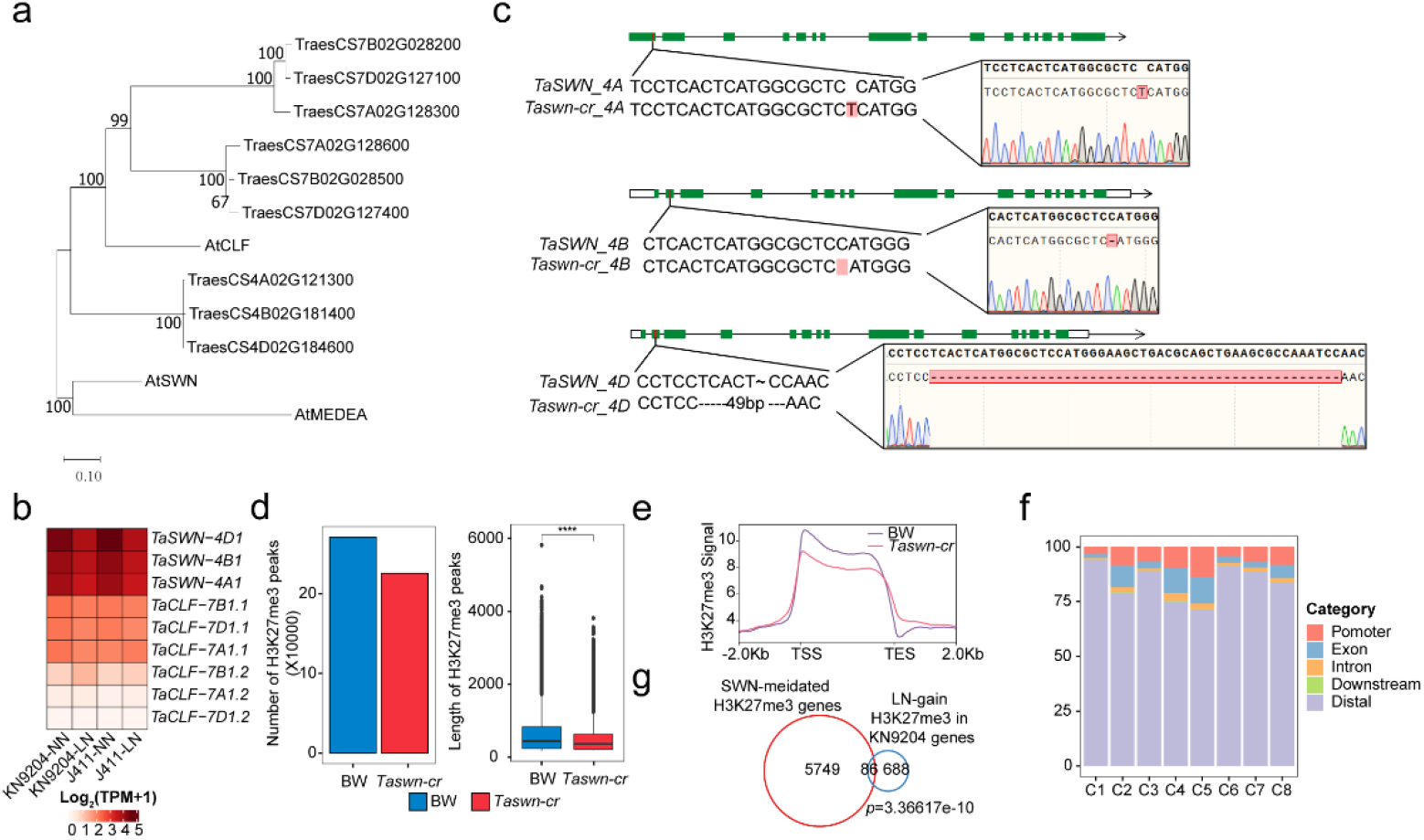
The generation and influence to global H3K27me3 level of *Taswn-cr*. a. A phylogenetic tree showing the evolutionary relationships between Ez proteins (one part of PRC2) from *Arabidopsis* and wheat. b. Relative expression of *TaSWN* and *TaCLF* genes under LN and NN conditions in KN9204 and J411. c. DNA sequence identification the mutated target sites in the three *TaSWN* genes in the *Taswn-cr* mutant. d. Peak numbers and lengths of H3K27me3 peaks in *Taswn-cr* and BW plants under different nitrogen conditions (Wilcox test: ns: *p*>0.05; *: *p*≤0.05; **: *p*≤0.01; ***: *p*≤0.001; ****: *p*≤0.0001). e. Profiles of H3K27me3 levels in *Taswn-cr* and BW under normal nitrogen conditions. f. Peak distribution of the differential H3K27me3 peaks in Fig. 6a. g. Overlap between genes marked by TaSWN-dependent H3K27me3 and genes marked by gain-H3K27me3 under LN condition in KN9204 (Fisher’s exact test was used to calculate the *p*-values for the overlaps).

